# Escape from antimicrobial CRISPR-Cas9 in *E. coli* ST131 depends on the genetic context of the target gene

**DOI:** 10.1101/2025.08.04.668416

**Authors:** Clàudia Morros-Bernaus, Joseph Westley, Ethan R. Wyrsch, Steven P. Djordjevic, Lihong Zhang, Anne FC. Leonard, William H. Gaze, David Sünderhauf, Stineke van Houte

## Abstract

*Escherichia coli* (*E. coli*) is a common bacterium in the human gut and an important cause of intestinal and extraintestinal infections. Some *E. coli* sequence types (ST) are associated with high pathogenicity. The Extraintestinal Pathogenic *E. coli* (ExPEC) ST131 is a globally distributed multidrug-resistant human pathogen associated with urinary tract and bloodstream infections. Antibiotic-resistant infections often lead to antibiotic treatment failure, underscoring the need of developing alternative treatments. The highly selective antimicrobial potential of CRISPR-Cas9 has been demonstrated in a range of model organisms. However, the effectiveness of CRISPR-Cas9 in combating ST131-associated infections and the consequences of CRISPR-Cas9 treatment, such as the emergence of escapers, remains unclear.

Here, we investigated the antimicrobial activity of CRISPR-Cas9 against ST131 and assessed the frequency and genetic basis of escape. We conjugatively delivered CRISPR-Cas9 to ST131 isolates which carried cefotaxime-resistance-encoding target gene *bla*_CTX-M-15_ in the chromosome and characterized escape subpopulations. Two main types of escapers emerged: *bla*_CTX-M-15_-positive escapers carried dysfunctional CRISPR-Cas9 systems and arose at a ∼10^−5^ frequency. Instead, *bla*_CTX-M-15_-negative escapers presented chromosomal deletions involving *bla*_CTX-M-15_ loss. The frequency of *bla*_CTX-M-15_ loss depended on the *bla*_CTX-M-15_ genetic context. Specifically, *bla*_CTX-M-15_-negative escapers emerged at low frequency (∼10^−5^) in isolates where *bla*_CTX-M-15_ was located downstream of insertion sequence (IS) IS*Ecp1*, while escapers emerged with high frequency (∼10^−3^) in isolates where *bla*_CTX-M-15_ was flanked by IS*26*. This work emphasizes how the genetic context of target genes can drive the outcome of CRISPR-Cas9 tools, where the presence of IS*26* may drive increased frequencies of escape.

**IMPORTANCE:** In the past decade CRISPR-Cas9 has emerged as an efficient antimicrobial tool capable of selective elimination of targeted bacteria. Even though it has been well described that bacteria can evolve to escape targeting by CRISPR-Cas9, the mechanisms of bacterial escape and their consequences remain largely elusive. In this study, we demonstrate the antimicrobial efficacy of CRISPR-Cas9 against natural isolates of *Escherichia coli* ST131, a clinically relevant pathogen, and elucidate the mechanism of escape from antimicrobial activity. We identify two distinct mechanisms of escape, which involve either dysfunctional CRISPR-Cas9 activity, or loss of the target gene (*bla*_CTX-M-15_), with the latter occurring at frequencies that depend on the genetic context of the target gene. These findings provide important insights into the frequency and mechanisms of bacterial escape from CRISPR-Cas9-based antimicrobials and offer a foundation for the development of more effective treatments.

## INTRODUCTION

Antimicrobial resistance (AMR) is a global health threat. By 2050, up to 8.22 million annual deaths are predicted to be associated with AMR (1). The misuse and overuse of antibiotics has drastically accelerated the emergence of AMR (2), which contributes to antibiotic treatment failure (3). In this context, *Escherichia coli* is one of the most important pathogens, accounting for the highest number of AMR-attributable deaths in 2019 (4). Therefore, it is recognized by the World Health Organization (WHO) as a priority pathogen for which urgent development of new antimicrobials is needed (5).

Extraintestinal Pathogenic *E. coli* (ExPEC) strains are among the most common Gram-negative pathogens in humans (6). They are associated with numerous types of infections, including urinary tract infections (UTIs), which can develop into bloodstream infections (7–9). Since 2000, Sequence Type (ST) ST131 is the most common pandemic lineage in the clinic (8). The ST131 lineage represents multidrug-resistant pathogens frequently associated with extended-spectrum beta-lactamases (ESBL), aminoglycoside and fluoroquinolone resistance, and several virulence factors (7,9–11). CTX-M enzymes are among the most prevalent type of ESBL (12,13), due to their global dissemination (12,14). The genes encoding these enzymes, *bla*_CTX-M_ genes, are found in both chromosomes and plasmids (14,15), often as part of highly mobile genetic structures, surrounded by insertion sequences (IS), transposons and integrons (14). These mobile genetic elements (MGEs) not only contribute to AMR mobilisation but can also act as promoter sequences, regulating expression levels of surrounding genes (14,16,17). Of these, IS are the smallest self-mobilizing units (0.7-2.5 kB) that typically code for a single transposase (Tnp), which allows their mobilisation (18,19). Several families of ISs, notably IS*Ecp1* and IS*26*, are widely distributed across *E. coli* ST131 genomes and are frequently located within or flanking CTX-M gene structures (20–22). While IS*Ecp1* is commonly found as a single copy upstream of the AMR gene (14), IS*26* is often found in multiple copies of the same orientation that flank the AMR gene, in what is known as a pseudo-compound transposon (PCT) (23,24).

ST131-associated infections are difficult to treat. In fact, higher antibiotic treatment failure rates have been reported compared to non-ST131 infections (25,26). In an urgent need to find alternative treatments, CRISPR-Cas9 can be used as a promising novel antimicrobial (27,28). In natural populations, CRISPR-Cas systems act as a prokaryotic immune system against MGE infections. Since its discovery CRISPR-Cas9 in particular has been developed as a highly sequence-specificity tool that can recognize a short (20 bp) specific DNA sequence and subsequently cleave it (29). The tool combines a single-guide RNA (*sgRNA*) with a Cas9 nuclease; the *sgRNA* recognises the target sequence while the Cas9 nuclease executes target cleavage by introducing a blunt-ended double-strand DNA break (DSB) into the target genome (30). DSBs are strong genotoxic lesions that lead to cell death (31). This highly specific antimicrobial effect of CRISPR-Cas9 has been studied extensively over the past ten years (32–36). However, the consequences of CRISPR-Cas9 treatment failure and the long-term effects of its use remain largely unexplored.

Here, we assessed the antimicrobial potential of CRISPR-Cas9 targeting *bla*_CTX-M-15_, the most reported ESBL gene among *E. coli* ST131 (13). We used a modified version of the broad-host range conjugative plasmid pKJK5::csg (37) to deliver CRISPR-Cas9, and programmed CRISPR-Cas9 to target four different *E. coli* ST131 isolates derived from human stool samples (38), all of them carrying a chromosomal copy of *bla*_CTX-M-15_. While strong antimicrobial activity was observed, we found escapers of CRISPR-Cas9 targeting for all isolates. We observed two main types of escapers, either presenting dysfunctional CRISPR-Cas9, or chromosomal rearrangements leading to loss of the target gene. Interestingly, for the second type of escapers, we found a direct impact of the *bla*_CTX-M-15_ genetic context (either surrounded by IS*Ecp1* or IS*26*) on the escape frequencies, emphasizing the importance of the genetic context of a target gene on escape from CRISPR-Cas9.

## RESULTS

### CRISPR-Cas9 targeting of a chromosomally encoded *bla*_CTX-M-15_ gene in human-associated *E*. *coli* ST131 isolates causes high levels of sequence-specific killing

We used the broad-host range conjugative plasmid pKJK5 as a delivery vector of a CRISPR-Cas9 cassette (csgc: *cas9, sgRNA, gfp* and *catB*), either targeting *bla*_CTX-M-15_ (pKJK5::csgc[*bla*_CTX-M-15_]) or a non-targeting control (pKJK5::csgc[NT]). We first verified successful delivery of pKJK5::csgc into a recipient *E. coli* DH5α lacking *bla*_CTX-M-15_, and found no significant differences (p = 0.68) between treatment and control (***Figure 1***), showing that the delivery efficiency of pKJK5::csgc is independent of the *sgRNA* target and that the system can be acquired in the absence of a CRISPR-Cas9 target.

**Figure 1:**
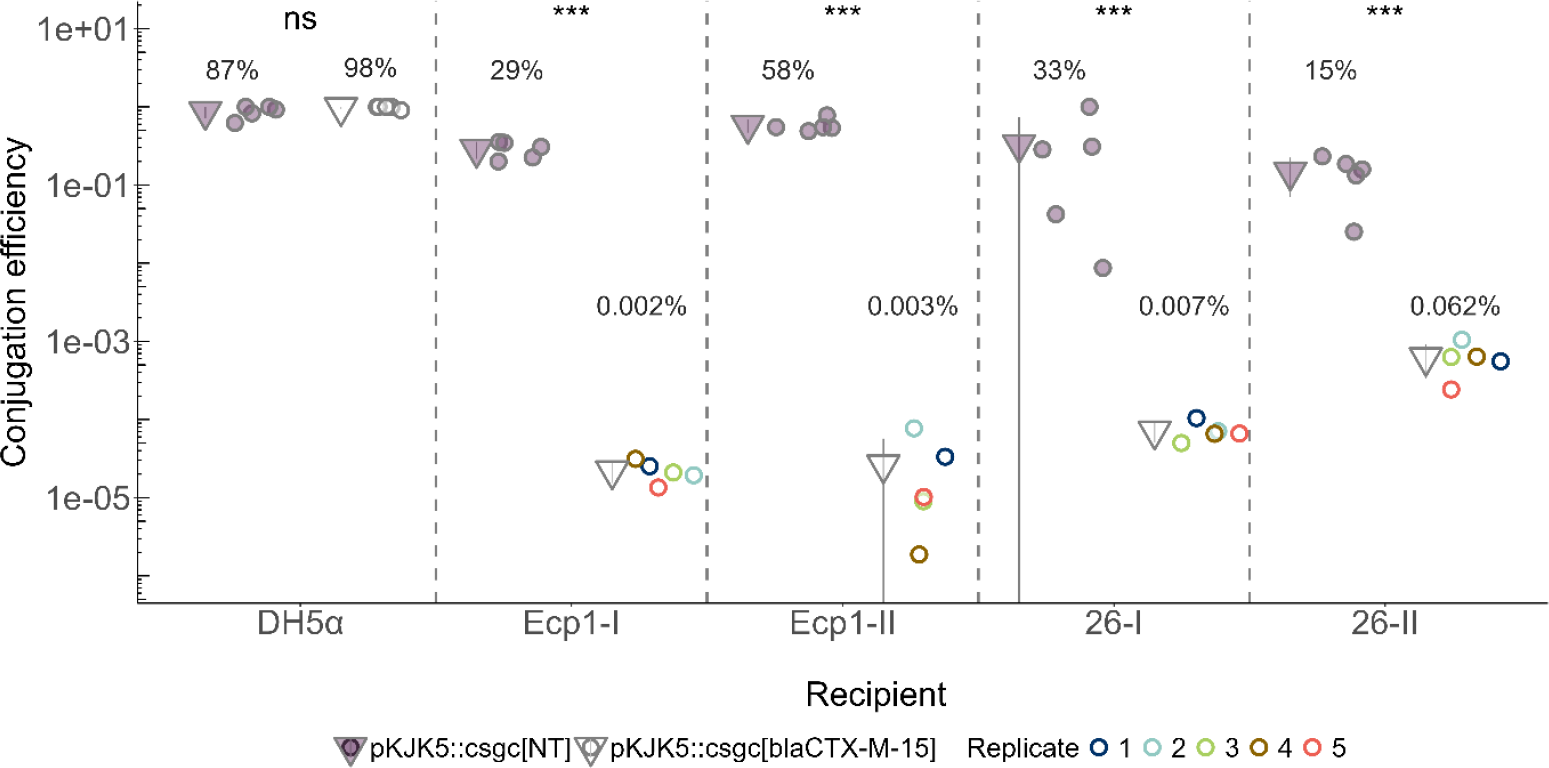
Conjugation efficiencies for either pKJK5::csgc[NT] (purple) or pKJK5::csgc[*bla*_CTX-M-15_] (white) across different recipients: either *E. coli* DH5α, lacking *bla*_CTX-M-15_, or the respective *E. coli* ST131 isolates, all with a chromosomally encoded *bla*_CTX-M-15_. Conjugation efficiencies were calculated dividing the number of transconjugants by the number of recipients. The conjugation efficiencies from the five replicates are represented as circles and the colour represents the replicate number. The distinct colours assigned to each replicate in the pKJK5::csgc[*bla*_CTX-M-15_] treatment facilitates tracking the replicate origin of the escapers throughout the manuscript. The means of the conjugation efficiencies are represented as triangles ± standard deviation (bars). p-values <0.001 (***), non-significant (ns).

Next, we assessed CRISPR-Cas9 targeting activity in four *E. coli* ST131 isolates (Ecp1-I, Ecp1-II, 26-I and 26-II); which all carry a single chromosomally encoded *bla*_CTX-M-15_ copy. Completed genome sequences were generated for all four isolates to enable us to have an unambiguous assessment of *bla*_CTX-M-15_ location and copy number. Across isolates, pKJK5::csgc[NT] achieved significantly higher conjugation efficiencies (ranging from 15 ± 8% to 58 ± 11%)) than pKJK5::csgc[*bla*_CTX-M-15_] (ranging from 0.002 ± 0.0007% to 0.062 ± 0.03%; p<0.001) (***Figure 1***). This difference was attributed to the non-viability of transconjugants when CRISPR-Cas9 targets the chromosomally encoded *bla*_CTX-M-15_.

While strong CRISPR-Cas9 antimicrobial activity was demonstrated, all isolates contained subpopulations of *E. coli* ST131 transconjugants (escapers) able to survive acquisition of pKJK5::csgc[*bla*_CTX-M-15_].

Overall escape frequencies varied across isolates ((4.53 ± 5.30) x 10^−5^ to (4.23 ± 1.95) x 10^−3^) (***Figure 2A***). To better understand the genetic basis of escape and their relative contribution to the overall escape frequencies, we determined cefotaxime resistance phenotypes for each of the escapers, since resistance to cefotaxime is known to be conferred by CTX-M enzymes (15,39). This showed that both cefotaxime-resistant and cefotaxime-sensitive escapers were found across isolates (***Supplementary figure 3***), revealing that *E. coli* ST131 could escape from CRISPR-Cas9 while either maintaining or losing the target gene.

**Figure 2:**
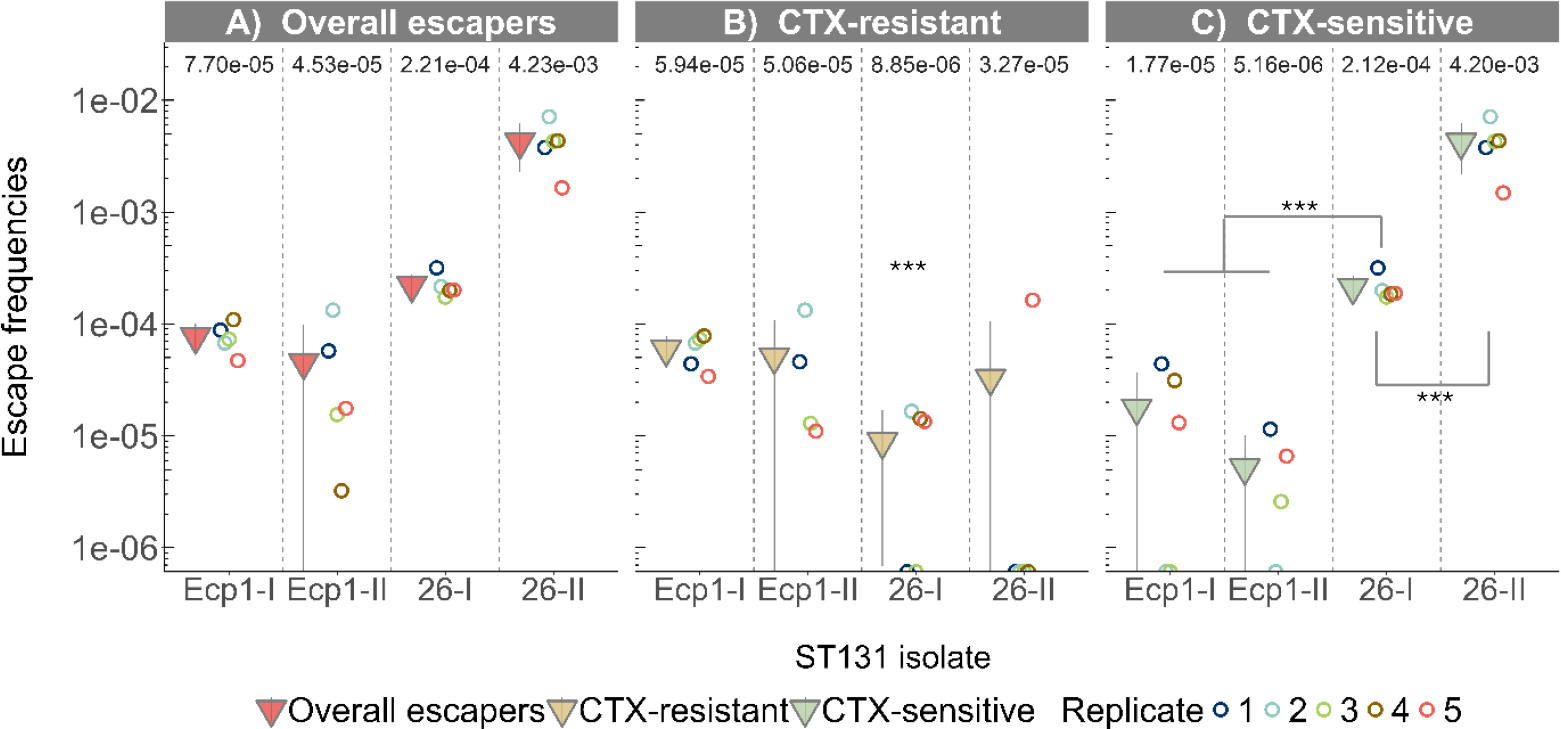
Mean escape frequencies from the CRISPR-Cas9 treatment for the four *E. coli* ST131 isolates. **A**) Overall escape frequencies independent of the cefotaxime resistance phenotype. These frequencies were calculated as the relative conjugation efficiency between the conjugation efficiency of pKJK5::csgc[*bla*_CTX-M-15_] for each replicate and the average conjugation efficiency of pKJK5::csgc[NT]. **B-C**) Cefotaxime-resistant (**B**) and -sensitive (**C**) escape frequencies. The escape frequencies for each phenotype were determined as the relative conjugation efficiency between the cefotaxime-resistant or cefotaxime-sensitive conjugation efficiencies of pKJK5::csgc[*bla*_CTX-M-15_] for each replicate and the average conjugation efficiency of pKJK5::csgc[NT]. The escape frequencies from the five replicates are represented as circles and the colour represents the replicate number. The mean escape frequencies, which include replicates with no escapers for a specific phenotype, are represented as triangles ± standard deviation (bars). Replicate 4 from isolate Ecp1-II was excluded from cefotaxime-resistant and cefotaxime-sensitive escape frequencies as only one escaper could be directly recovered from the filter mating assay. p-values <0.001 (***).

**Figure 3:**
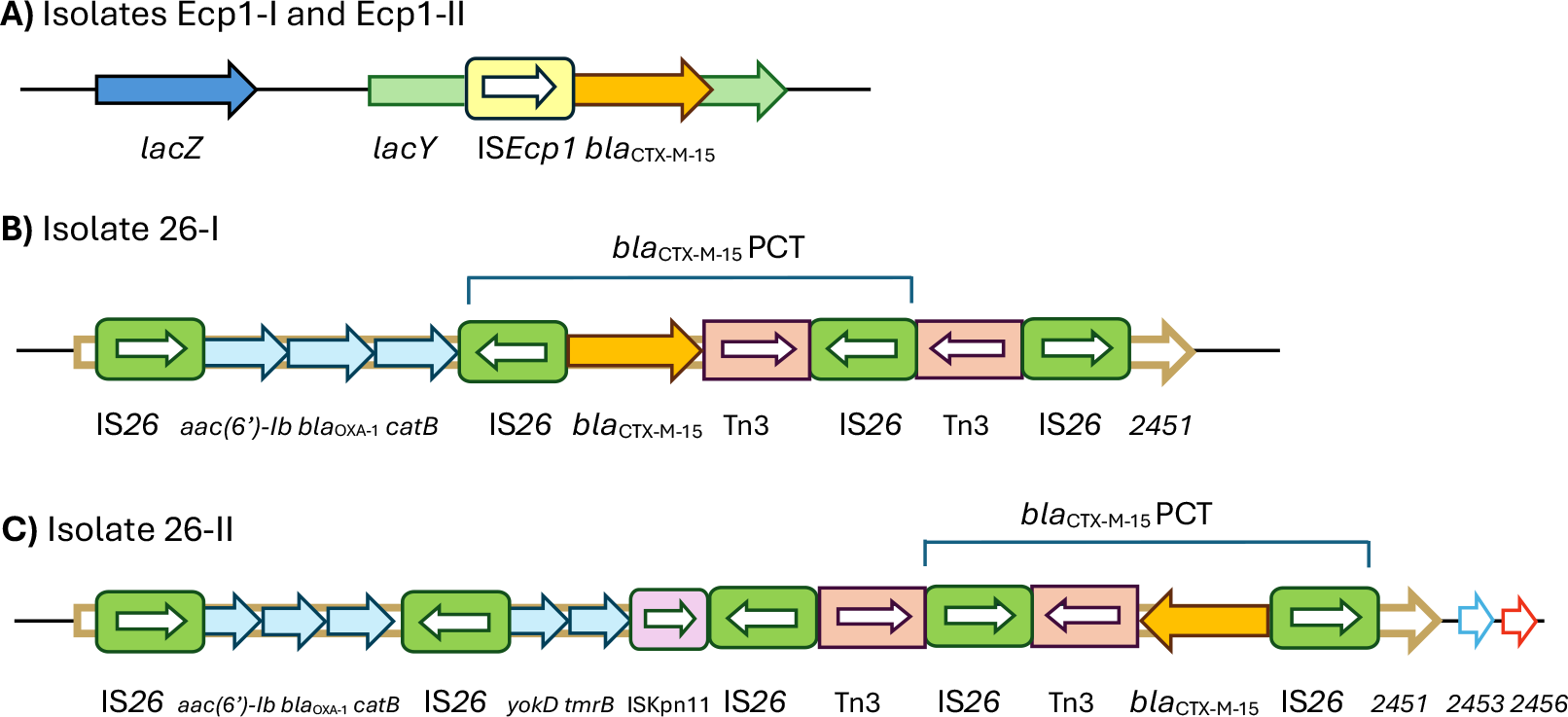
Genetic context of *bla*_*CTX-M-15*_ (orange arrow) for the four *E. coli* ST131 isolates, ORFs and gene distances not drawn to scale. **A**) Isolate Ecp1-I and Ecp1-II share the same ISEcp1 genetic context disrupting a *lacY* gene. The *lacZ* gene is found 1.2 kB upstream of the IS*Ecp1*. IS*Ecp1* is represented as a yellow arrow box. **B-C**) IS*26* genetic contexts for isolates 26-I (**B**) and 26-II (**C**), with a *bla*_CTX-M-15_ pseudo-compound transposon (PCT). For both isolates, the respective overall IS*26*-contained sequences are found disrupting the gene with unknow function (*EC958_2451*), shown in the figure as 2451. The two Tn3, shown as pink boxes, represent a single Tn3 split into two truncated ORFs, likely due to IS*26* disruption that led to an inversion of one of the parts. IS*26* is represented as a green arrow box. *2453* and *2456* represent the genes *EC958_2453 and EC958_2456*.

### Cefotaxime-resistant escapers have dysfunctional CRISPR-Cas9 systems

Cefotaxime-resistant escape frequencies were generally low (∼10^−5^) but showed significant variation across isolates (***Figure 2B*;** p<0.001). Post-hoc testing revealed this was due to isolate 26-I having a significantly lower escape frequency than all others (p<0.001). Nevertheless, the escape frequencies for isolates 26-I and 26-II should be interpreted cautiously, as only few escapers were detected, and their presence was inconsistent across replicates (***Figure 2B*** and ***Supplementary figure 3***). Therefore, in-depth analysis of cefotaxime-resistant escapers was only performed for isolates Ecp1-I and Ecp1-II. First, we sequenced the *bla*_CTX-M-15_ gene to genetically confirm the cefotaxime-resistant phenotypes and found no mutations in the target site or PAM. We then hypothesized that escapers likely survived CRISPR-Cas9 targeting through acquiring mutations in *cas9* and/or *sgRNA* (csgc) on the pKJK5::csgc[*bla*_CTX-M-15_]. To address this, we assessed CRISPR-Cas9 integrity using PCR. All transconjugants from the pKJK5::csgc[NT] control showed expected *bla*_CTX-M-15_ and csgc amplicons, but deletions, duplications, and partial or complete lack of amplification within *cas9* and/or *sgRNA* were found for 48% (Ecp1-I) and 64% (Ecp1-II) of the escapers (***Supplementary figure 4A***). Furthermore, one Ecp1-II escaper exhibited an IS150 transposition into *cas9*. While IS150 is present in both the donor (K-12 MG1655) and the recipient (ST131 isolate Ecp1-II) genome, the presence of a single nucleotide polymorphism unique for IS150 in the K-12 genome suggested that the transposition into pKJK5::csg[*bla*_CTX-M-15_] occurred in the donor prior to its delivery to Ecp1-II. The different mutations were distributed across the *cas9* and *sgRNA* sequence (***Supplementary figure 4B-C***), with no obvious patterns revealed except for a hotspot in the α-helical lobe found in escapers from isolate Ecp1-I (***Supplementary figure 4B***) (40).

Those escapers that did not exhibit identifiable csgc mutations were subjected to a phenotypic CRISPR-Cas9 functionality assay. We mated escapers with a donor bacterium bearing either targeted plasmid pCTX15 (carrying the wildtype *bla*_CTX-M-15_) or untargeted plasmid pCTRL (***Supplementary figure 5A***) and calculated their relative conjugation efficiencies (pCTRL/pCTX15). Cefotaxime-sensitive escapers lacking *bla*_CTX-M-15_ were used as potential positive CRISPR-Cas9 controls. For both Ecp1-I and Ecp1-II most cefotaxime-sensitive escapers demonstrated protection against pCTX15 (***Supplementary figure 5B-C***). In contrast, no protection was observed for cefotaxime-resistant escapers (***Supplementary figure 5B-C***), indicating dysfunctional CRISPR-Cas9 activity. Only one cefotaxime-resistant escaper (5.10 Ecp1-II) showed a relative conjugation efficiency for all the replicates compatible with functional CRISPR-Cas9 activity (***Supplementary figure 5C***), which could indicate coexistence of the wildtype *bla*_CTX-M-15_ and a functional pKJK5::csgc[*bla*_CTX-M-15_]. This escaper showed no mutations in *bla*_CTX-M-15_, and short-read WGS did not reveal relevant mutations in pKJK5::csgc[*bla*_CTX-M-15_] that could interfere with CRISPR-Cas9 activity. Therefore, coexistence could be attributed to an unknown mechanism.

To summarize, we found that cefotaxime-resistant escape frequencies were ∼10^−5^ across isolates and all the escapers carried *bla*_CTX-M-15_ with no mutations in the protospacer or PAM sequence. Using genotypic and phenotypic screening, we attributed cell survival to dysfunctional CRISPR-Cas9 systems.

### Cefotaxime-sensitive escapers lost *bla*_*CTX-M-15*_ in a manner dependent on its genetic context

To understand the differential cefotaxime-sensitive escape frequencies found across isolates (***Figure 2C***), we sought to understand the genetic context of *bla*_CTX-M-15_. Long-read genome analyses of the ancestral untreated isolates revealed three different genetic contexts. In isolates Ecp1-I and Ecp1-II, *bla*_CTX-M-15_ is found downstream of an insertion sequence (IS) IS*Ecp1*, which disrupts *lacY* in the *lac* operon (***Figure 3A***). In contrast, isolates 26-I and 26-II carry *bla*_CTX-M-15_ flanked by two IS*26* copies of the same orientation, in a pseudo-compound transposon (PCT) (18,23). This *bla*_CTX-M-15_ PCT is found surrounded by either two or three additional IS*26* copies respectively, creating two slightly distinct overall IS*26*-contained sequences. These sequences share the same genome location in both isolates, disrupting a gene of unknown function (*EC958_2451*) (***Figure 3B-C***). Significant differences were found between cefotaxime-sensitive escape frequencies of the three genetic contexts (p-values for each combination < 0.001). Isolates with an IS*Ecp1* genetic context showed significantly lower cefotaxime-sensitive escape frequencies than isolate 26-I (4-copy IS*26* context), which in turn had a significantly lower escape frequency than isolate 26-II (5-copy IS*26* context). Overall, this indicates a significantly higher escape frequency through loss of *bla*_CTX-M-15_ in isolates with an IS*26* genetic context compared to IS*Ecp1*. Consequently, different approaches were employed to characterize the escapers based on their genetic context.

### An IS*Ecp1* genetic context is associated with large-scale genomic deletions alongside *bla*_CTX-M-15_ loss

For cefotaxime-sensitive escapers from Ecp1-I (n=19) and Ecp1-II (n=19) (***Supplementary figure 3***), PCR and subsequent Sanger sequencing of the amplicons revealed complete loss of IS*Ecp1* + *bla*_CTX-M-15_ for most of the escapers (n=36) and partial deletions including the protospacer sequence (the 20 bp targeted by pKJK5::csgc[*bla*_CTX-M-15_]) for the remaining escapers (n=2) (***Supplementary figure 6A***).

To better characterize the dimensions of the IS*Ecp1* + *bla*_CTX-M-15_ deletions, a β-galactosidase assay was performed, taking advantage of the proximal *lacZ* gene (***Figure 3A***). This enzymatic assay allows phenotypic screening of blue or white colonies based on presence or absence, respectively, of β-galactosidase, the enzyme encoded by *lacZ*. A negative phenotype (white) indicated deletions of IS*Ecp1* + *bla*_CTX-M-15_ together with *lacZ* (***Supplementary figure 6B***). In contrast, a positive phenotype (blue) delimited the start of the deletion within 1.2 kB (somewhere in between *lacZ* and IS*Ecp1*) (***Supplementary figure 6C***). Both phenotypes were observed across escapers, with β-galactosidase-negative escapers being more abundant (n=30) (***Supplementary figure 6D***). Short-read WGS was performed on a representative escaper from each phenotype and isolate to confirm their genotypic basis.

Additionally, all cefotaxime-sensitive escapers from Ecp1-I and Ecp1-II were subjected to both the csgc genotypic and phenotypic CRISPR-Cas9 functionality assay previously described. All escapers showed csgc amplicons of expected lengths. In the phenotypic assay most showed functional CRISPR-Cas9 systems (n=32). Nevertheless, a small subset showed dysfunctional CRISPR-Cas9 activity (n=6), highlighting that both the loss of *bla*_CTX-M-15_ and the presence of a dysfunctional system can co-occur (***Supplementary figure 5B-C***).

### An IS*26* genetic context is associated with small-scale genomic deletions of *bla*_CTX-M-15_ facilitated by the presence of two flanking IS*26* copies

For isolates 26-I and 26-II, both with an IS*26* genetic context, 26-II was chosen as a representative for further study due to its higher IS*26* load (***Figure 3 B-C***). First, hybrid WGS was performed for a subset of escapers (n=10) which revealed either the deletion of the *bla*_CTX-M-15_ PCT (n=8) or larger deletions including downstream chromosomal sequences (n=2). All the deletions led to a single remaining IS*26* chromosomal copy from the two original ones.

We used these representative data to inform PCRs assaying the deletion of the *bla*_CTX-M-15_ PCT (***Supplementary figure 7A***) in all the cefotaxime-sensitive escapers (n=98). The ancestral isolate and the cefotaxime-resistant escapers (used as positive controls) showed the expected 4.9 kB amplicon covering the *bla*_CTX-M-15_ PCT alongside a smaller amplicon, likely a PCR artefact resulting from recombination between the multiple copies of IS*26*. Crucially, cefotaxime-sensitive escapers (n=94) lacked the 4.9 kB amplicon and only displayed a 972 bp amplicon, which corresponded to a single IS*26* copy in agreement with the deletion of the *bla*_CTX-M-15_ PCT observed in the WGS (***Supplementary figure 7B***). Additionally, 3 escapers, including the two with larger deletions characterized by WGS, showed no amplicon, agreeing with the presence of larger deletions thus avoiding primer binding (***Supplementary figure 7A and C-D***). Finally, one escaper showed a <4.9 kb amplicon, likely explained by a partial deletion within *bla*_CTX-M-15_, thereby conferring loss of the resistant phenotype and escape of CRISPR-Cas9 targeting.

To summarize, the emergence of cefotaxime-sensitive escapers was significantly impacted by the genetic context of *bla*_CTX-M-15._ While all cefotaxime-sensitive escapers exhibited chromosomal rearrangements involving the loss of *bla*_CTX-M-15_, an IS*Ecp1* genetic context was associated with lower escape frequencies and large-scale deletions. In contrast, an IS*26* genetic context revealed higher escape frequencies, primarily driven by small-scale deletions (*bla*_CTX-M-15_ PCT) occurring between the two flanking IS*26* elements.

## DISCUSSION

Here, we demonstrated the sequence-specific antimicrobial activity of CRISPR-Cas9 by targeting the chromosomally encoded ESBL gene *bla*_CTX-M-15_ in human-associated isolates of the *E. coli* lineage ST131. The delivery of a *bla*_CTX-M-15_ targeting CRISPR-Cas9 cassette using the broad host-range conjugative plasmid pKJK5::csgc achieved a significant reduction (from 236-fold to 22.000-fold, depending on the isolate) in conjugation efficiency compared to a non-targeting control, explained by the non-viability of *E. coli* ST131 transconjugants. This demonstrates the antimicrobial use of CRISPR-Cas9 against *E. coli* ST131 isolates, which can be found in antimicrobial resistant UTIs, for which non-antibiotic treatments are urgently needed (41). Promisingly, the use of pKJK5::csgc is compatible with probiotics, which are already prophylactic and therapeutic treatment options for recurrent UTIs (42–44). However, we observed widespread escape of CRISPR-Cas9 targeting across isolates and identified both escapers that retained the *bla*_CTX-M-15_ gene and carried dysfunctional CRISPR-Cas9, and escapers that lost *bla*_CTX-M-15_ through chromosomal rearrangements. Crucially, for the escapers with *bla*_CTX-M-15_ deletions, we found significantly different escape frequencies depending on the genetic context of the target gene, with increased escape frequencies (up to 813-fold) when *bla*_CTX-M-15_ is found surrounded by two copies of IS*26*, as part of an IS*26* pseudo-compound transposon (PCT), compared to when it is found downstream of an IS*Ecp1*.

Cefotaxime-resistant (*bla*_CTX-M-15_*-*positive) escapers retained an intact version of *bla*_CTX-M-15_, suggesting that CRISPR-Cas9 escape through mutations in the protospacer (the 20 bp targeted by pKJK5::csgc[*bla*_CTX-M-15_]) or protospacer adjacent motif (PAM) sequence is rare in these isolates. As mutations in the target site were previously reported for other genes (36,45–49), we hypothesize that the absence of such escapers in our setup may be attributed to mutations in *bla*_CTX-M-15_ occurring below the detection limit of the assay (∼10^−6^). Instead, all cefotaxime-resistant escapers presented dysfunctional CRISPR-Cas9 systems. In line with previous literature (35,45,46,48,50,51), we found deletions, duplications and insertions across *cas9* and *sgRNA*. Interestingly, we only found one escaper with an IS transposition into *cas9*, even though this is a commonly reported event (36,46,47,50,52). To minimize escapers with dysfunctional CRISPR-Cas9 systems, several solutions have been proposed, including the use of multi-target arrays (35,36,48), the manipulation of the CRISPR-Cas9 plasmid copy number (50), coupling CRISPR-Cas antimicrobials with CRISPR-regulated toxin-antitoxin systems (ATTACK) (53) or the use of non-DNA based delivery strategies such as nano-sized CRISPR complexes (54), especially as plasmid-based delivery relies on often heterogenous transcriptional activity of recipients (55). However, the optimal solution to this problem has yet to be established.

In contrast, cefotaxime-sensitive escapers revealed *bla*_CTX-M-15_ deletions, and this loss of the target gene occurred in significantly higher frequencies (up to 813-fold) for isolates with an IS*26* genetic context compared to an IS*Ecp1* genetic context. The observed *bla*_CTX-M-15_ deletions were likely driven by homologous recombination (HR), which could either occur in the pre-existing population leading to positively selected genotypes upon CRISPR-Cas9 exposure, or as a DNA-damage repair mechanism after CRISPR-Cas9 targeting and a subsequent triggering of the SOS response (33,36,56). Escapers with successful DNA-damage repair could be associated with weaker CRISPR-Cas9 activity (33), poor *sgRNA* folding (57) or expression (58) and/or variability in the host DNA damage tolerance and responses (59). Deletions of *bla*_CTX-M-15_ compatible with HR were observed across isolates. For isolates Ecp1-I and Ecp1-II, sequencing of few cefotaxime-sensitive escapers revealed deletions with border homology (9-11 bp), likely indicating HR. In isolate 26-II, the most common deletion observed among cefotaxime-sensitive escapers (a deletion of the *bla*_CTX-M-15_ PCT leaving a single IS*26* copy in the chromosome; ***Supplementary figure 7B***), is consistent with the product of HR between the two directly oriented IS*26* copies (60–62) (***Supplementary figure 8A-B***). Recombination before CRISPR-Cas9 cleavage would release a circular molecule carrying *bla*_CTX-M-15_, known as a translocatable unit (TU) (18,24,60,63,64), (***Supplementary figure 8A***) which is expected to be readily lost in absence of self-replicative features (60) or through active targeting by CRISPR-Cas9. CRISPR-Cas9 cleavage of intermediate circular molecules from similar excised MGEs has been recently reported (65). Overall, the higher escape frequencies observed in an IS*26* genetic context could be attributed to the homology found between the several IS*26* copies (***Figure 3 B-C***), which may facilitate HR. In fact, HR between homologous IS elements is a common driver of chromosomal rearrangements in *E. coli* (66).

Alternatively, *bla*_CTX-M-15_ loss could be driven by an incomplete IS-mediated *bla*_CTX-M-15_ mobilization. Evidence of this was found in escapers from isolate 26-II with larger *bla*_CTX-M-15_ deletions lacking homology in the borders (***Supplementary figure 7C-D***). These deletions likely resulted from a Tnp26-dependent intramolecular copy-in mobilisation (24,67), in where the downstream genes *EC958_2453* and *EC958_2456* would have been used as respective intramolecular targets to mediate chromosomal excision, releasing a TU carrying the enclosed sequence (60) (***Supplementary figure 8CI-II***), which would be readily lost or targeted by CRISPR-Cas9. This mobilization pattern could also contribute to escapers with a *bla*_CTX-M-15_ PCT deletion, if using the sequence found downstream of the PCT as an intramolecular target site (***Supplementary figure 8CIII***). In contrast, we could not find evidence of *bla*_CTX-M-15_ deletions mediated by IS*Ecp1* mobilisation, as IS*Ecp1* mobilises together with downstream genes (18,68–71). Therefore, this would only be possible in *lacZ-*positive cefotaxime-sensitive escapers, which are a minority (***Supplementary figure 6D***). Sequencing of two *lacZ*-positive escapers revealed *bla*_CTX-M-15_ deletions incompatible with IS*Ecp1* mobilisation, suggesting that this is not a common escape mechanism.

To summarize, even though our data does not allow us to discern the exact escape mechanism that leads to chromosomal rearrangements, deletions resulting from both HR and incomplete IS-mediated *bla*_CTX-M-15_ mobilization are not mutually exclusive, and it is likely that the total escape population is a combinatorial result of these events, each occurring at different frequencies.

Escapers with chromosomal rearrangements involving the deletion of the target gene have been reported for other genes and bacterial species (33,36,49,57,72–74), including deletions with border homology, suggesting HR (33,75,76). Additionally, some studies have shown a reduction in escape frequencies when knocking out (33) or inhibiting (57) expression of RecA, an important protein involved in HR (77) and found in all the ST131 isolates used in this work. Interestingly, studies where the target gene is found within a chromosomally integrated MGE reported escapers in which the entire MGE was deleted, similar to the deletions of the *bla*_CTX-M-15_ PCT reported here. This occurred when the target genes were in a genomic island flanked by two directly oriented IS1193 copies (73), in several pathogenicity islands (49,78) and in prophages (78).

The presence of escapers with chromosomal rearrangements after a CRISPR-Cas9 treatment might be especially relevant when occurring at high frequencies, as for isolate 26-II (∼10^−3^). The emergence of this type of escapers could likely be reduced by choosing target genes independent from chromosomally integrated MGEs and with genetic contexts that present limited or no homology. In this study, escapers with *bla*_CTX-M-15_ loss were resensitized to cefotaxime, which could be a beneficial outcome for antibiotic therapy. However, this type of chromosomal rearrangements can also involve loss of other genes, leading to phenotypic changes (73,76). Furthermore, deletions resulting from successful DNA repair after CRISPR-Cas9 cleavage through HR could be accompanied by mobilisation of other HR-mediated MGEs (18). This might be especially relevant when using CRISPR-Cas9 against AMR-carrying bacteria, which often carry MGEs (18). While we did not directly observe this, such far-reaching genomic consequences of CRISPR-Cas9 targeting should also be kept in mind when using CRISPR-Cas9 as an editing tool, which often relies on HR (31,58).

Altogether, this study showed the antimicrobial activity of CRISPR-Cas9 against the chromosomally encoded AMR gene *bla*_CTX-M-15_ from human-associated *E. coli* ST131 isolates and identified escapers resulting from either dysfunctional CRISPR-Cas9 or with chromosomal rearrangements that led to the deletion of *bla*_CTX-M-15_. Our work showed the impact that the genetic context of the target gene has on escape frequencies, where an association with IS*26* led to a high frequency of *bla*_CTX-M-15_ loss. This research underlines the importance of understanding the genetic environment to be able to predict the treatment outcome of CRISPR-Cas9 antimicrobials and CRISPR-Cas9 gene editing. Finally, we also want to highlight the potential use of CRISPR-Cas9 as a tool to better characterize IS mobilization patterns by studying escapers arising from the targeting of the IS elements or cargo genes.

## METHODS

### Growth conditions, buffers and media

Lysogeny broth (LB), either liquid or mixed with agar, was used as growth media. Antibiotics were used to ensure plasmid maintenance, select for chromosomal markers or perform phenotypic assays at: chloramphenicol (Cm) 25 μg mL^-1^, gentamicin (Gm), kanamycin (Km) and streptomycin (Sm) all 50 μg mL^-1^, and cefotaxime (CTX) 5 μg mL^-1^. Glycerol stocks were prepared at 20% (w/v) and frozen at -70 °C. Sterile 0.9 % (w/v) NaCl was used as a buffer as indicated. Unless otherwise specified, all kits and reagents were used following manufacturer’s instructions and incubations were performed overnight (O/N) at 37 °C, 180 rpm.

### Bacterial strains

We selected the *E. coli* ST131 isolates used in this study (Ecp1-I, Ecp1-II, 26-I and 26-II) based on the genetic context of *bla*_CTX-M-15_ (genomes deposited on Genbank under BioProject PRJNA1281408). All the ST131 isolates belong to clade C, the most relevant in the clinic (79). Isolates Ecp1-I, Ecp1-II and 26-I were chromosomally tagged with an *aacC1* gene, conferring Gm resistance, through electroporation of a Tn5 transposon plasmid pBAMD1-6 (80), followed by PCR screening to confirm absence of residual plasmid as well as competition experiments against the respective ancestral isolates to verify that tagging did not incur a fitness cost. In brief, the modified strains were grown together with their wildtype variants, and colony counts revealed similar growth. Isolate 26-II was not genetically modified due to intrinsic Gm resistance. *E. coli* K-12 MG1655::*mCherry* (81) was used as CRISPR-Cas9 donor strain because it represses expression of *gfp* from pKJK5::csgc, which allows us to verify successful plasmid conjugation to the ST131 isolates (presence of transconjugants) by checking *gfp* expression. Further bacteria and plasmid information can be found in ***Supplementary table 1*** and ***2***, respectively.

### Development of the CRISPR-Cas9 system

Two CRISPR-Cas9 cassettes were designed with distinct *sgRNA* targets. The [*bla*_CTX-M-15_] *sgRNA* targets [CGCGTGATACCACTTCACCT]. This sequence is found within *bla*_CTX-M-15_ and followed by a protospacer adjacent motif (PAM) in the *E. coli* ST131 reference genome (82) and the genome of the four *E. coli* ST131 isolates used in the study. Furthermore, the sequence showed low predicted CRISPR-off target activity in both CRISPOR (83) and Cas-OFFinder (84). The non-targeting [NT] control *sgRNA* targets random nucleotide sequence [GGTAAGACCATTAGAAGTAG], 20 bp which we confirmed to be absent from all *E. coli* ST131 isolates. These cassettes were generated and transferred into pKJK5, resulting in pKJK5::csgc[*bla*_CTX-M-15_] or [NT], following protocols adopted from (37) (full details in ***Supplementary method 1*** and ***Supplementary table 3***).

### Filter mating conjugation *E. coli* K-12 MG1655 (donor) - *E. coli* ST131 (recipient)

Single colonies of donors (MG1655 pKJK5::csgc[*bla*_CTX-M-15_] or [NT]) and recipients (*E. coli* DH5α::SmR or the respective *E. coli* ST131 isolates) were grown O/N in 5 mL LB. Donors were supplemented with Cm to avoid segregational loss of pKJK5. Cells were washed twice in 5 mL NaCl, followed by OD600 adjustment to 0.5 – 0.6. Recipients were diluted 100-fold in NaCl. Filter mating was performed in a Millipore 1225 Sampling Manifold using a sterile Whatman Cyclopore Clear 0.2 µm 25mm polycarbonate membrane on top of a sterile Whatman glass microfiber filter, binder free, grade GF/C, 25mm. The vacuum manifold was sterilised using ethanol and UV before and between batches. Filters were washed by pumping through 2 mL of NaCl. Straight after, 1 mL of NaCl, 1 mL of donor and 1 mL of a 100-fold diluted recipient were pumped through. Five biological replicates were performed for each donor-recipient combination. Additionally, controls with donor-only, recipient-only and NaCl-only were performed and yielded results consistent with expectations, supporting the validity of the experiment. The polycarbonate filters were placed onto 10 % LB-agar plates and incubated for 48 h at 37 °C in the absence of antibiotics. Filters were recovered in 3 mL NaCl and vortexed. From the cell suspension, cells were recovered in LB, after which differential selective plating on LB agar was used to quantify the proportion of 1) donors (no antibiotic selection; donors identified by assessing *mCherry* expression using a stereo fluorescent lamp (Nightsea), 2) recipients (Gm) and 3) transconjugants (Gm + Cm). Glycerol stocks were made with the remaining cell suspensions.

For each replicate, conjugation efficiency was calculated by dividing transconjugant concentrations (CFU/mL) by recipient concentrations (CFU/mL) and means were determined for each isolate and treatment. For each replicate from the pKJK5::csgc[*bla*_CTX-M-15_] treatment, escape frequencies were calculated as a relative conjugation efficiency by dividing the conjugation efficiency of pKJK5::csgc[*bla*_CTX-M-15_] for each replicate with the average conjugation efficiency of pKJK5::csgc[NT]. The mean escape frequency per isolate was calculated as the average of all five biological replicates.

### Recovery of *E. coli* ST131 escapers (*E. coli* ST131 pKJK5::csgc[*bla*_*CTX-M-15*_]) and phenotypic analysis

*E. coli* ST131 pKJK5::csgc[*bla*_CTX-M-15_] escapers were recovered on Gm + Cm plates and the presence of *bla*_*CTX-M-15*_ was verified based on their cefotaxime resistance profile. To do so, individual clones were replated onto LB agar containing (i) Cm + CTX and (ii) Cm to assess the cefotaxime phenotype while ensuring CRISPR-Cas9 plasmid maintenance. Additionally, we generated glycerol stocks of escapers after O/N incubation in LB + Cm. The same procedure was used to select for 100 transconjugants from the non-targeting treatment (*E. coli* ST131 pKJK5::csgc[NT]). A visual representation of the filter mating assay and the phenotypic screen of the escapers can be found in ***Supplementary figure 1***.

To calculate cefotaxime-resistant and cefotaxime-sensitive escape frequencies, the cefotaxime resistance profiles were assessed for all escapers recovered directly from the filter mating assay for isolates Ecp1-I, Ecp1-II and 26-I. For isolate 26-II, due to the larger size of the escape population recovered, ten escapers per replicate were randomly selected for resistance profiling. The proportions of each cefotaxime phenotype were then used to estimate cefotaxime-resistant and cefotaxime-sensitive transconjugants concentrations (CFU/mL) per replicate. Conjugation efficiencies for both phenotypes were calculated by dividing the respective transconjugant concentrations by the recipient concentrations for each replicate of the pKJK5::csgc[*bla*_CTX-M-15_] treatment. Finally, cefotaxime-resistant and cefotaxime-sensitive escape frequencies were calculated as a relative conjugation efficiency by dividing the cefotaxime-resistant or cefotaxime-sensitive conjugation efficiencies of pKJK5::csgc[*bla*_CTX-M-15_] for each replicate with the average conjugation efficiency of pKJK5::csgc[NT]. The mean escape frequency per phenotype and isolate was calculated as the average of all biological replicates, including those where no cefotaxime-resistant or cefotaxime-sensitive escapers were found. Replicate 4 from isolate Ecp1-II was excluded from the calculations of cefotaxime-resistant and cefotaxime-sensitive escape frequencies as only one escaper was directly recovered from the filter mating assay.

### Genotypic analysis of *bla*_**CTX-M-15**_

PCR was used to study *bla*_CTX-M-15_ presence in escapers. To obtain enough DNA template material, frozen glycerol stocks of escapers were scraped with a pipette tip and the tip was swirled in 10 μL water to suspend attached material. The suspension was then heated to 95 °C for 15 minutes and centrifugated at 3.500 rpm for 5 minutes. For cefotaxime-resistant and cefotaxime-sensitive escapers from isolates Ecp1-I and Ecp1-II, *bla*_CTX-M-15_ presence was studied using primers with binding sites within the gene (***Supplementary figure 2A***). Furthermore, for the cefotaxime-sensitive escapers primers designed to amplify IS*Ecp1* + *bla*_CTX-M-15_ were also used (***Supplementary figure 2A***). Phusion High-Fidelity polymerase (Thermo Scientific) was used in both PCRs. Additionally, ExoSAP-cleaned (NEB) PCR amplicons were Sanger sequenced. In cefotaxime-sensitive escapers from isolate 26-II, the deletion of the *bla*_CTX-M-15_ PCT was studied using primers annealing outside the PCT (***Supplementary figure 2B***) with 1x VeriFi Hot Start Polymerase (PCR Biosystems). Across PCRs, when no amplicon was found, 16S PCR or amplification of other genes were used to verify template presence. Moreover, the respective ancestral isolates and transconjugants from the non-targeting control (pKJK5::csgc[NT]) were used as positive controls for the presence of *bla*_CTX-M-15_. All primers can be found in ***Supplementary table 3***.

### Genotypic analysis of CRISPR-Cas9 integrity

Cefotaxime-resistant and cefotaxime-sensitive escapers from isolates Ecp1-I and Ecp1-II were subjected to a genotypic CRISPR-Cas9 integrity assay to understand whether mutations had occurred in *cas9* and/or the *sgRNA*. Four sets of primers generating overlapping amplicons were used, including a region upstream of *cas9* and downstream of the *sgRNA*. Additionally, a fifth set was used to specifically amplify the *sgRNA* (***Supplementary figure 2C***). PCRs were performed using 2x PCRBIO Taq Mix Red (PCR Biosystems). Amplicons with unexpected lengths (i.e. amplicons with an estimated length >60 bp different from the expected amplicon length) were purified from the agarose gel using the Monarch® DNA Gel Extraction Kit Protocol (NEB) and Sanger sequenced. Transconjugants from the pKJK5::csgc[NT] control and the ancestral isolates were used as controls, and 16S amplification was used to verify template presence. DNA template for the PCRs was obtained as described above. All primers can be found in ***Supplementary table 3***.

### Phenotypic CRISPR-Cas9 functionality assay

To check for the presence of dysfunctional CRISPR-Cas9 activity in cefotaxime-resistant escapers from isolates Ecp1-I and Ecp1-II that showed expected lengths of csgc amplicons and, in cefotaxime-sensitive escapers, a phenotypic CRISPR-Cas9 functionality assay was performed. For this assay, liquid conjugations were performed with donors harbouring either plasmid pCTX15, which we engineered to carry *bla*_CTX-M-15_, or plasmid pCTRL, which we engineered to carry a mutated version of *bla*_CTX-M-15_ that allows escape from CRISPR-Cas9 targeting through a silent point mutation of the PAM sequence. Specific details about plasmid engineering and the assay can be found in ***Supplementary method 1*** and ***2***.

### Phenotypic β-galactosidase assay

To characterize the IS*Ecp1 bla*_CTX-M-15_ deletions found in cefotaxime-sensitive escapers from isolates Ecp1-I and Ecp1-II, a phenotypic β-galactosidase assay was performed that makes use of the proximity of *bla*_CTX-M-15_ to the *lacZ* gene. Individual escapers were plated onto 0.2 μg mL^-1^ X-Gal LB-agar plates. Colonies were screened for white (β-galactosidase / *lacZ*-negative) or blue (β-galactosidase / *lacZ*-positive) phenotypes. Cefotaxime-resistant escapers, transconjugants from the pKJK5::csgc[NT] control and the ancestral isolates were used as positive controls.

### Sequencing

Whole genome sequencing was performed to obtain genomic details of the four *E. coli* ST131 ancestral isolates and several escapers from the CRISPR-Cas9 treatment. For the ancestral isolates, DNA extraction was performed from LB + CTX 3 μg mL^-1^ ON cultures using FastDNA Spin Kit (MP Biomedicals). The extracted DNA was treated with 2 μL of RNAseA 20 mg mL^-1^ for 10 minutes at 37 °C and purified with SPRISelect beads (Beckman Coulter). Sequencing libraries were prepared with 2 μg of purified DNA using the ONT Ligation Sequencing Kit (Oxford Nanopore Technologies) and long-read sequencing was performed using PromethION (Exeter Sequencing Facility). Genomes were assembled using *Unicycler* (version 0.5.0) (85), with automated annotations generated using *RASTtk* (86). Short-read WGS (MicrobesNG, Birmingham) was performed for one β-galactosidase-positive and one β-galactosidase-negative cefotaxime-sensitive escaper from isolates Ecp1-I and Ecp1-II. Similarly, short-read WGS (MicrobesNG, Birmingham) was also performed for escaper 5.10 from isolate Ecp1-II. Hybrid WGS (MicrobesNG, Birmingham) was performed for ten cefotaxime-sensitive escapers from isolate 26-II (two per each filter mating replicate) and for one transconjugant from the pKJK5::csgc[NT] control. Genome assembly was performed using *Flye* (87) and the respective deletions were characterized. Benchling was used for DNA visualization (88).

### Data analysis

Data processing, visualisation and statistical analyses were performed in R version 4.4.2 using R studio 2023.9.1.494 (89). Statistical modelling was performed using package *lme4* (version 1.1-37) (90). Data processing and plotting were performed using packages *tidyverse* (version 2.0.0) (91), *ggplot2* (version 3.5.1) (92), *gggenes* (version 0.5.1) (93), *patchwork* (version 1.3.1) (94) and *grid* (version 4.4.2) (89). Significance of fixed effects was determined through comparing nested models using chi-squared tests (α = 0.05), beginning with a global model that included all biologically relevant fixed effects and interaction terms. Interaction term statistical significance was always tested first, and where interactions were statistically significant, all constituent fixed effects were retained in the model. R package *DHARMa* (version 0.4.6) (95) was used to diagnose potential model issues. Specifically, we checked if residuals deviation were deviated from the expected distribution, if the model had over or under dispersion, if the datasets contained highly influential outliers, and if the residuals were heteroscedastic. Post-hoc analyses were performed using *emmeans* (version 1.10.2) (96).

Conjugation efficiency of CRISPR-Cas9 and the cefotaxime-resistant and cefotaxime-sensitive escape frequencies were modelled using binomial family generalised linear models (GLMs) and a logit link function, with the number of colonies counted for the recipients and transconjugants as the weights value. Due to large differences in the count of transconjugants between treatments, concentrations of recipients and transconjugants (CFU/mL) were logarithmically transformed (when the dataset included zeros, log10+1 was used).

Our global model of conjugation efficiency of CRISPR-Cas9 included the predictor variables plasmid identity, isolate identity, and an interaction between plasmid and isolate identity. For the cefotaxime-resistant and cefotaxime-sensitive escape frequency analyses we modelled the predictor variable genetic context of the *bla*_CTX-M-15_. For the cefotaxime-sensitive escape frequencies, model validation revealed slight deviation in the first quartile of the residuals *vs* predicted plot. However, this model remained the most appropriate, and the large observed effect sizes mean it is improbable we are drawing erroneous conclusions from this analysis. For the cefotaxime-resistant escape frequencies, model validation revealed some influential outliers (replicates with no escapers) and model optimization was performed removing outliers.

For the phenotypic CRISPR-Cas9 functionality assay in *E. coli* K-12 MG1655 carrying either pKJK5::csgc[NT] or pKJK5::csgc[*bla*_CTX-M-15_] the relative conjugation efficiencies (pCTRL/pCTX15) were modelled using a Gamma family GLM with a log link function and with CRISPR-Cas9 plasmid as a predictor variable. For the phenotypic CRISPR-Cas9 functionality assay with escapers, the data were individually modelled for each isolate. For both datasets, pCTRL/pCTX15 was modelled using a Gamma family GLM with a log link function, with cefotaxime phenotype as a predictor variable. Model validation revealed some heteroscedasticity for isolate Ecp1-II, which could be attributed to the presence of few cefotaxime-sensitive escapers with dysfunctional CRISPR-Cas9 systems. Nevertheless, the Gamma GLM remained the most appropriate for our data.

## DATA AVAILABILITY

The genome of the *E. coli* ST131 isolates (Ecp1-I, Ecp1-II, 26-I, 26-II) were deposited in PRJNA1281408. The assembled genomes of escapers with *bla*_CTX-M-15_ deletions from isolates Ecp1-I, Ecp1-II and 26-II and for escaper 5.10 Ecp1-II can be found in *to be determined*. The data analysis can be found in GitHub *to be determined*. All the raw data and the full escape characterization can be found in the raw data excel file in *to be determined*. Additional experimental details, methods, tables and figures can be found in the online version of this article.

## ACKNOWLEDGEMENTS AND FUNDING

S.V.H. gratefully acknowledges funding from the Biotechnology and Biological Sciences Research council (BBSRC; BB/R010781/1), the Lister Institute for Preventative Medicine, and the Joint Programme Initiative AntiMicrobial Resistance (JPI-AMR) ‘Harissa’ call (MISTAR; MR/W031191/1)). D.S was supported in part by grant MR/N0137941/1 for the GW4 BIOMED MRC DTP, awarded to the Universities of Bath, Bristol, Cardiff and Exeter from the Medical Research Council (MRC)/UKRI. Genome sequencing of the ST131 isolates was supported by funding from the Natural Environment Research Council awarded to AFCL (award number NE/R013748/1) and utilised equipment funded by the UK Medical Research Council (MRC) Clinical Research Infrastructure Initiative (award number MR/M008924/1). For the purpose of open access, the author has applied a ‘Creative Commons Attribution (CC BY) licence to any Author Accepted Manuscript version arising from this submission.

## Notes

### Competing Interest Statement

The authors have declared no competing interest.

### Summary of Updates

The manuscript has been updated to add the supplementary data which was lacking before.

## Bibliography

1. Naghavi M, Vollset SE, Ikuta KS, Swetschinski LR, Gray AP, Wool EE, et al. Global burden of bacterial antimicrobial resistance 1990–2021: a systematic analysis with forecasts to 2050. The Lancet. 2024 Sep 28;404(10459):1199–226.

2. Dadgostar P. Antimicrobial Resistance: Implications and Costs. Infect Drug Resist. 2019 Dec 20;Volume 12:3903–10.

3. Haney EF, Hancock REW. Addressing Antibiotic Failure—Beyond Genetically Encoded Antimicrobial Resistance. Front Drug Discov. 2022;2:892975.

4. Murray CJ, Ikuta KS, Sharara F, Swetschinski L, Robles Aguilar G, Gray A, et al. Global burden of bacterial antimicrobial resistance in 2019: a systematic analysis. Lancet. 2022;399(10325):629–55.

5. World Health Organization. Prioritization of pathogens to guide discovery, research and development of new antibiotics for drug-resistant bacterial infections, including tuberculosis. 2017.

6. Poolman JT, Wacker M. Extraintestinal Pathogenic Escherichia coli, a Common Human Pathogen: Challenges for Vaccine Development and Progress in the Field. J Infect Dis. 2016 Jan 1;213(1):6–13.

7. Daga AP, Koga VL, Soncini JGM, de Matos CM, Perugini MRE, Pelisson M, et al. Escherichia coli Bloodstream Infections in Patients at a University Hospital: Virulence Factors and Clinical Characteristics. Front Cell Infect Microbiol. 2019 Jun 6;9:191.

8. Manges AR, Geum HM, Guo A, Edens TJ, Fibke CD, Pitout JDD. Global Extraintestinal Pathogenic Escherichia coli (ExPEC) Lineages. Clin Microbiol Rev. 2019 Jun 12;32(3):e00135–18.

9. Rogers BA, Sidjabat HE, Paterson DL. Escherichia coli O25b-ST131: a pandemic, multiresistant, community-associated strain. J Antimicrob Chemother. 2011 Jan;66(1):1–14.

10. Johnson JR, Johnston B, Clabots C, Kuskowski MA, Castanheira M. Escherichia coli sequence type ST131 as the major cause of serious multidrug-resistant E. coli infections in the United States. Clin Infect Dis. 2010 Aug 1;51(3):286–94.

11. Pitout JDD, Finn TJ. The evolutionary puzzle of Escherichia coli ST131. Infect Genet Evol. 2020 Jul;81:104265.

12. Cantón R, Coque TM. The CTX-M beta-lactamase pandemic. Curr Opin Microbiol. 2006 Oct;9(5):466–75.

13. Nicolas-Chanoine MH, Bertrand X, Madec JY. Escherichia coli ST131, an intriguing clonal group. Clin Microbiol Rev. 2014 Jul;27(3):543–74.

14. Cantón R, González-Alba JM, Galán JC. CTX-M Enzymes: Origin and Diffusion. Front Microbiol. 2012 Apr 2;3:110.

15. Rossolini GM, D’Andrea MM, Mugnaioli C. The spread of CTX-M-type extended-spectrum beta-lactamases. Clin Microbiol Infect. 2008 Jan 1;14(Suppl 1):33–41.

16. Siguier P, Gourbeyre E, Chandler M. Bacterial insertion sequences: their genomic impact and diversity. FEMS Microbiol Rev. 2014 Sep;38(5):865–91.

17. Zhao WH, Hu ZQ. Epidemiology and genetics of CTX-M extended-spectrum β-lactamases in Gram-negative bacteria. Crit Rev Microbiol. 2013 Feb;39(1):79–101.

18. Partridge SR, Kwong SM, Firth N, Jensen SO. Mobile Genetic Elements Associated with Antimicrobial Resistance. Clin Microbiol Rev. 2018 Aug 1;31(4):e00088–17.

19. Mahillon J, Chandler M. Insertion sequences. Microbiol Mol Biol Rev. 1998 Sep;62(3):725–74.

20. Pitout JDD, Chen L. The Significance of Epidemic Plasmids in the Success of Multidrug-Resistant Drug Pandemic Extraintestinal Pathogenic Escherichia coli. Infect Dis Ther. 2023 Apr;12(4):1029–41.

21. Woodford N, Carattoli A, Karisik E, Underwood A, Ellington MJ, Livermore DM. Complete nucleotide sequences of plasmids pEK204, pEK499, and pEK516, encoding CTX-M enzymes in three major Escherichia coli lineages from the United Kingdom, all belonging to the international O25:H4-ST131 clone. Antimicrob Agents Chemother. 2009 Oct;53(10):4472–82.

22. Pajand O, Rahimi H, Darabi N, Roudi S, Ghassemi K, Aarestrup FM, et al. Arrangements of Mobile Genetic Elements among Virotype E Subpopulation of Escherichia coli Sequence Type 131 Strains with High Antimicrobial Resistance and Virulence Gene Content. mSphere. 2021 Aug 25;6(4):e0055021.

23. Varani A, He S, Siguier P, Ross K, Chandler M. The IS6 family, a clinically important group of insertion sequences including IS26. Mob DNA. 2021 Mar;23;12(1):11.

24. Harmer CJ, Hall RM. IS26 and the IS26 family: versatile resistance gene movers and genome reorganizers. Microbiol Mol Biol Rev. 2024 Jun 27;88(2):e0011922.

25. Can F, Azap OK, Seref C, Ispir P, Arslan H, Ergonul O. Emerging Escherichia coli O25b/ST131 clone predicts treatment failure in urinary tract infections. Clin Infect Dis. 2015 Feb 15;60(4):523–7.

26. Syre H, Hetland MAK, Bernhoff E, Bollestad M, Grude N, Simonsen GS, et al. Microbial risk factors for treatment failure of pivmecillinam in community-acquired urinary tract infections caused by ESBL-producing Escherichia coli. APMIS. 2020 Mar;128(3):232–41.

27. Pursey E, Sünderhauf D, Gaze WH, Westra ER, van Houte S. CRISPR-Cas antimicrobials: Challenges and future prospects. PLoS Pathog. 2018 Jun 1;14(6):e1006990.

28. Mayorga-Ramos A, Zúñiga-Miranda J, Carrera-Pacheco SE, Barba-Ostria C, Guamán LP. CRISPR-Cas-Based Antimicrobials: Design, Challenges, and Bacterial Mechanisms of Resistance. ACS Infect Dis. 2023 Jul 14;9(7):1283–302.

29. Jinek M, Chylinski K, Fonfara I, Hauer M, Doudna JA, Charpentier E. A programmable dual-RNA-guided DNA endonuclease in adaptive bacterial immunity. Science (1979). 2012 Aug 17;337(6096):816–21.

30. Jiang F, Doudna JA. CRISPR-Cas9 Structures and Mechanisms. Annu Rev Biophys. 2017 May 22;46:505–29.

31. Xue C, Greene EC. DNA Repair Pathway Choices in CRISPR-Cas9-Mediated Genome Editing. Trends Genet. 2021 Jul;37(7):639–56.

32. Citorik RJ, Mimee M, Lu TK. Sequence-specific antimicrobials using efficiently delivered RNA-guided nucleases. Nat Biotechnol. 2014 Nov;32(11):1141–5.

33. Cui L, Bikard D. Consequences of Cas9 cleavage in the chromosome of Escherichia coli. Nucleic Acids Res. 2016 May 19;44(9):4243–51.

34. Bikard D, Euler CW, Jiang W, Nussenzweig PM, Goldberg GW, Duportet X, et al. Exploiting CRISPR-Cas nucleases to produce sequence-specific antimicrobials. Nat Biotechnol. 2014 Nov;32(11):1146–50.

35. Uribe R V., Rathmer C, Jahn LJ, Ellabaan MMH, Li SS, Sommer MOA. Bacterial resistance to CRISPR-Cas antimicrobials. Sci Rep. 2021 Dec 1;11(1):17267.

36. Reuter A, Hilpert C, Dedieu-Berne A, Lematre S, Gueguen E, Launay G, et al. Targeted-antibacterial-plasmids (TAPs) combining conjugation and CRISPR/Cas systems achieve strain-specific antibacterial activity. Nucleic Acids Res. 2021 Apr 6;49(6):3584–98.

37. Sünderhauf D, Klümper U, Pursey E, Westra ER, Gaze WH, van Houte S. Removal of AMR plasmids using a mobile, broad host-range CRISPR-Cas9 delivery tool. Microbiology (Reading). 2023 May;169(5):001334.

38. Leonard AFC, Zhang L, Balfour AJ, Garside R, Hawkey PM, Murray AK, et al. Exposure to and colonisation by antibiotic-resistant E. coli in UK coastal water users: Environmental surveillance, exposure assessment, and epidemiological study (Beach Bum Survey). Environ Int. 2018 May;114:326–33.

39. Paterson DL, Bonomo RA. Extended-Spectrum β-Lactamases: a Clinical Update. Clin Microbiol Rev. 2005 Oct;18(4):657–86.

40. Jinek M, Jiang F, Taylor DW, Sternberg SH, Kaya E, Ma E, et al. Structures of Cas9 Endonucleases Reveal RNA-Mediated Conformational Activation. Science (1979). 2014 Mar 14;343(6176):1247997–1247997.

41. Sihra N, Goodman A, Zakri R, Sahai A, Malde S. Nonantibiotic prevention and management of recurrent urinary tract infection. Nat Rev Urol. 2018 Dec;15(12):750–76.

42. Durrani B, Mohammad A, Ljubetic BM, Dobberfuhl AD. The Potential Role of Persister Cells in Urinary Tract Infections. Curr Urol Rep. 2023 Nov;24(11):541–51.

43. Reid G, Bruce AW. Probiotics to prevent urinary tract infections: the rationale and evidence. World J Urol. 2006 Feb;24(1):28–32.

44. Gupta V, Nag D, Garg P. Recurrent Urinary Tract Infections in Women: How Promising is the Use of Probiotics? Indian J Med Microbiol. 2017 Jul 1;35(3):347–54.

45. Jiang W, Bikard D, Cox D, Zhang F, Marraffini LA. RNA-guided editing of bacterial genomes using CRISPR-Cas systems. Nat Biotechnol. 2013 Jan 29;31(3):233–9.

46. Neil K, Allard N, Roy P, Grenier F, Menendez A, Burrus V, et al. High-efficiency delivery of CRISPR-Cas9 by engineered probiotics enables precise microbiome editing. Mol Syst Biol. 2021 Oct;17(10):e10335.

47. Hamilton TA, Pellegrino GM, Therrien JA, Ham DT, Bartlett PC, Karas BJ, et al. Efficient inter-species conjugative transfer of a CRISPR nuclease for targeted bacterial killing. Nat Commun. 2019 Oct 4;10(1):4544.

48. Gomaa AA, Klumpe HE, Luo ML, Selle K, Barrangou R, Beisel CL. Programmable removal of bacterial strains by use of genome-targeting CRISPR-cas systems. mBio. 2014 Jan 28;5(1):e00928–13.

49. Vercoe RB, Chang JT, Dy RL, Taylor C, Gristwood T, Clulow JS, et al. Cytotoxic Chromosomal Targeting by CRISPR/Cas Systems Can Reshape Bacterial Genomes and Expel or Remodel Pathogenicity Islands. PLoS Genet. 2013 Apr;9(4):e1003454.

50. Li Q, Sun M, Lv L, Zuo Y, Zhang S, Zhang Y, et al. Improving the Editing Efficiency of CRISPR-Cas9 by Reducing the Generation of Escapers Based on the Surviving Mechanism. ACS Synth Biol. 2023 Mar 17;12(3):672–80.

51. Aparicio T, de Lorenzo V, Martínez-García E. CRISPR/Cas9-Based Counterselection Boosts Recombineering Efficiency in Pseudomonas putida. Biotechnol J. 2018 May;13(5):e1700161.

52. Sheng Y, Wang H, Ou Y, Wu Y, Ding W, Tao M, et al. Insertion sequence transposition inactivates CRISPR-Cas immunity. Nat Commun. 2023 Jul 20;14(1):4366.

53. Wang R, Shu X, Zhao H, Xue Q, Liu C, Wu A, et al. Associate toxin-antitoxin with CRISPR-Cas to kill multidrug-resistant pathogens. Nat Commun. 2023 Apr 12;14(1):2078.

54. Kang YK, Kwon K, Ryu JS, Lee HN, Park C, Chung HJ. Nonviral Genome Editing Based on a Polymer-Derivatized CRISPR Nanocomplex for Targeting Bacterial Pathogens and Antibiotic Resistance. Bioconjug Chem. 2017 Apr 19;28(4):957–67.

55. Cyriaque V, Ibarra-Chávez R, Kuchina A, Seelig G, Nesme J, Madsen JS. Single-cell RNA sequencing reveals plasmid constrains bacterial population heterogeneity and identifies a non-conjugating subpopulation. Nat Commun. 2024;15:5853.

56. Moreb EA, Hoover B, Yaseen A, Valyasevi N, Roecker Z, Menacho-Melgar R, et al. Managing the SOS Response for Enhanced CRISPR-Cas-Based Recombineering in E. coli through Transient Inhibition of Host RecA Activity. ACS Synth Biol. 2017 Dec 15;6(12):2209–18.

57. Vialetto E, Miele S, Goren MG, Yu J, Yu Y, Collias D, et al. Systematic interrogation of CRISPR antimicrobials in Klebsiella pneumoniae reveals nuclease-, guide-and strain-dependent features influencing antimicrobial activity. Nucleic Acids Res. 2024 Jun 10;52(10):6079–91.

58. Collias D, Vialetto E, Yu J, Co K, Almási É d. H, Rüttiger AS, et al. Systematically attenuating DNA targeting enables CRISPR-driven editing in bacteria. Nat Commun. 2023 Feb 8;14(1):680.

59. Moreb EA, Hoover B, Yaseen A, Valyasevi N, Roecker Z, Menacho-Melgar R, et al. Managing the SOS Response for Enhanced CRISPR-Cas-Based Recombineering in E. coli through Transient Inhibition of Host RecA Activity. ACS Synth Biol. 2017 Dec 15;6(12):2209–18.

60. Harmer CJ, Hall RM. IS26-Mediated Formation of Transposons Carrying Antibiotic Resistance Genes. mSphere. 2016 Apr 6;1(2):e00038–16.

61. Harmer CJ, Hall RM. IS26-Mediated Precise Excision of the IS26-aphA1a Translocatable Unit. mBio. 2015 Dec 8;6(6):e01866–15.

62. Aihara M, Gotoh Y, Shirahama S, Matsushima Y, Uchiumi T, Kang D, et al. Generation and maintenance of the circularized multimeric IS26-associated translocatable unit encoding multidrug resistance. Commun Biol. 2024;7:597.

63. Harmer CJ, Moran RA, Hall RM. Movement of IS26-associated antibiotic resistance genes occurs via a translocatable unit that includes a single IS26 and preferentially inserts adjacent to another IS26. mBio. 2014 Oct 7;5(5):e01801–14.

64. Harmer CJ, Moran RA, Hall RM. Movement of IS26-Associated Antibiotic Resistance Genes Occurs via a Translocatable Unit That Includes a Single IS26 and Preferentially Inserts Adjacent to Another IS26. mBio. 2014 Oct 7;5(5).

65. Wang P, Du X, Zhao Y, Wang W, Cai T, Tang K, et al. Combining CRISPR/Cas9 and natural excision for the precise and complete removal of mobile genetic elements in bacteria. Appl Environ Microbiol. 2024 Apr 1;90(4):e0009524.

66. Raeside C, Gaffé J, Deatherage DE, Tenaillon O, Briska AM, Ptashkin RN, et al. Large chromosomal rearrangements during a long-term evolution experiment with Escherichia coli. mBio. 2014 Sep 9;5(5):e01377–14.

67. He S, Hickman AB, Varani AM, Siguier P, Chandler M, Dekker JP, et al. Insertion Sequence IS26 Reorganizes Plasmids in Clinically Isolated Multidrug-Resistant Bacteria by Replicative Transposition. mBio. 2015 Jun 9;6(3):e00762.

68. Poirel L, Decousser JW, Nordmann P. Insertion Sequence ISEcp1B Is Involved in Expression and Mobilization of a blaCTX-M β-Lactamase Gene. Antimicrob Agents Chemother. 2003 Sep;47(9):2938–45.

69. Zong Z, Partridge SR, Iredell JR. ISEcp1-Mediated Transposition and Homologous Recombination Can Explain the Context of blaCTX-M-62 Linked to qnrB2. Antimicrob Agents Chemother. 2010 Jul;54(7):3039–42.

70. Poirel L, Lartigue MF, Decousser JW, Nordmann P. ISEcp1B-Mediated Transposition of blaCTX-M in Escherichia coli. Antimicrob Agents Chemother. 2005 Jan;49(1):447–50.

71. Wachino JI, Yamane K, Kimura K, Shibata N, Suzuki S, Ike Y, et al. Mode of Transposition and Expression of 16S rRNA Methyltransferase Gene rmtC Accompanied by ISEcp1. Antimicrob Agents Chemother. 2006 Sep;50(9):3212–5.

72. Qi LS, Larson MH, Gilbert LA, Doudna JA, Weissman JS, Arkin AP, et al. Repurposing CRISPR as an RNA-Guided Platform for Sequence-Specific Control of Gene Expression. Cell. 2013 Feb 28;152(5):1173–83.

73. Selle K, Klaenhammer TR, Barrangou R. CRISPR-based screening of genomic island excision events in bacteria. Proc Natl Acad Sci U S A. 2015 Jun 30;112(26):8076–81.

74. Guan J, Wang W, Sun B. Chromosomal Targeting by the Type III-A CRISPR-Cas System Can Reshape Genomes in Staphylococcus aureus. mSphere. 2017 Nov 15;2(6):e00403–17.

75. Lam KN, Spanogiannopoulos P, Soto-Perez P, Alexander M, Nalley MJ, Bisanz JE, et al. Phage-delivered CRISPR-Cas9 for strain-specific depletion and genomic deletions in the gut microbiome. Cell Rep. 2021 Nov 2;37(5):109930.

76. Cañez C, Selle K, Goh YJ, Barrangou R. Outcomes and characterization of chromosomal self-targeting by native CRISPR-Cas systems in Streptococcus thermophilus. FEMS Microbiol Lett. 2019 May 1;366(9):fnz105.

77. Hu J, Crickard JB. All who wander are not lost: the search for homology during homologous recombination. Biochem Soc Trans. 2024 Feb 28;52(1):367–77.

78. Wang P, Du X, Zhao Y, Wang W, Cai T, Tang K, et al. Combining CRISPR/Cas9 and natural excision for the precise and complete removal of mobile genetic elements in bacteria. Appl Environ Microbiol. 2024 Apr 17;90(4):e0009524.

79. Kocsis B, Gulyás D, Szabó D. Emergence and Dissemination of Extraintestinal Pathogenic High-Risk International Clones of Escherichia coli. Life (Basel). 2022 Dec 10;12(12):2077.

80. Martínez-García E, Aparicio T, de Lorenzo V, Nikel PI. New transposon tools tailored for metabolic engineering of Gram-negative microbial cell factories. Front Bioeng Biotechnol. 2014 Oct 28;2:46.

81. Klümper U, Riber L, Dechesne A, Sannazzarro A, Hansen LH, Sørensen SJ, et al. Broad host range plasmids can invade an unexpectedly diverse fraction of a soil bacterial community. ISME J. 2015;9(4):934–45.

82. Forde BM, Ben Zakour NL, Stanton-Cook M, Phan MD, Totsika M, Peters KM, et al. The Complete Genome Sequence of Escherichia coli EC958: A High Quality Reference Sequence for the Globally Disseminated Multidrug Resistant E. coli O25b:H4-ST131 Clone. PLoS One. 2014 Aug 15;9(8):e104400.

83. Concordet JP, Haeussler M. CRISPOR: intuitive guide selection for CRISPR/Cas9 genome editing experiments and screens. Nucleic Acids Res. 2018 Jul 2;46(W1):W242–5.

84. Bae S, Park J, Kim JS. Cas-OFFinder: a fast and versatile algorithm that searches for potential off-target sites of Cas9 RNA-guided endonucleases. Bioinformatics. 2014 May 15;30(10):1473–5.

85. Wick RR, Judd LM, Gorrie CL, Holt KE. Unicycler: Resolving bacterial genome assemblies from short and long sequencing reads. PLoS Comput Biol. 2017 Jun 1;13(6):e1005595.

86. Brettin T, Davis JJ, Disz T, Edwards RA, Gerdes S, Olsen GJ, et al. RASTtk: A modular and extensible implementation of the RAST algorithm for building custom annotation pipelines and annotating batches of genomes. Sci Rep. 2015 Feb 10;5:8365.

87. Kolmogorov M, Yuan J, Lin Y, Pevzner PA. Assembly of long, error-prone reads using repeat graphs. Nat Biotechnol. 2019 May;37(5):540–6.

88. [Biology Software]. Benchling [Internet]. 2024. Available from: https://benchling.com

89. Posit team. RStudio: Integrated Development Environment for R [Internet]. Boston, MA: Posit Software, PBC; 2023. Available from: http://www.posit.co/

90. Bates D, Mächler M, Bolker BM, Walker SC. Fitting Linear Mixed-Effects Models Using lme4. J Stat Softw [Internet]. 2015 Oct 7;67(1):1–48. Available from: https://www.jstatsoft.org/article/view/v067i01

91. Wickham H, Averick M, Bryan J, Chang W, D’ L, Mcgowan A, et al. Welcome to the Tidyverse. J Open Source Softw [Internet]. 2019 Nov 21;4(43):1686. Available from: https://joss.theoj.org/papers/10.21105/joss.01686

92. Hadley Wickham. ggplot2: Elegant Graphics for Data Analysis [Internet]. Springer-Verlag New York; 2016. Available from: https://ggplot2.tidyverse.org

93. David Wilkins. gggenes: Draw Gene Arrow Maps in “ggplot2” [Internet]. 2020. Available from: https://CRAN.R-project.org/package=gggenes

94. Thomas Lin Pedersen. patchwork: The Composer of Plots [Internet]. 2025. Available from: https://patchwork.data-imaginist.com

95. Hartig F. DHARMa: Residual Diagnostics for Hierarchical (Multi-Level / Mixed) Regression Models. CRAN: Contributed Packages. 2016.

96. Lenth R V. emmeans: Estimated Marginal Means, aka Least-Squares Means. CRAN: Contributed Packages. 2017.

