## Supplementary data for "Escape from antimicrobial CRISPR-Cas9 in *E. coli* ST131 depends on the genetic context of the target gene"

### SUPPLEMENTARY METHODS

#### *Supplementary method 1* – Design and engineering of plasmids

The respective CRISPR-Cas9 cassettes ([*bla*$_{CTX\text{-}M\text{-}15}$] and [NT]) were designed based on a previous version (1), following the same methodology except for the use of pACYCduet-1 as a carrier plasmid instead of pMARQ. Standard molecular cloning described in (1) was used to introduce the [*bla*$_{CTX\text{-}M\text{-}15}$] and [NT] *sgRNA* respectively into pACYC_csg and transfer them into pDOC, generating the respective pDOC::csg. In addition to *cas9*, *sgRNA* and *gfp* (green fluorescent protein), which were already present in the original CRISPR-Cas9 cassette (*csg*), a *catB* chloramphenicol resistance reporter gene was added, giving rise to the csgc cassette (*cas9*, *sgRNA, gfp* and *catB*). *catB* was added by PCR amplification from pACYCduet-1 with primers introducing restrictions sites for AgeI-HF (NEB) (***Supplementary table 3***), using Phusion High-Fidelity polymerase (Thermo Scientific). Ligations of the purified gene with the digested plasmids were performed using T4 DNA ligase (NEB), generating the respective pDOC_csgc plasmids. Finally, the cassettes, which were surrounded by homology arms matching the *intI1* region on pKJK5, were transferred onto pKJK5 using λ-red-assisted recombineering as described in (1), except for Cm being used to select for recombinants.

For the phenotypic CRISPR-Cas9 functionality assay pSEVA251(2) was used to generate pCTX15 and pCTRL. Plasmid pCTX15 carried a wildtype *bla*$_{CTX\text{-}M\text{-}15}$ gene targeted by a functional pKJK5::csgc[*bla*$_{CTX\text{-}M\text{-}15}$], while the plasmid pCTRL carried *bla*$_{CTX\text{-}M\text{-}15}$ with a silent point mutation (CGG to CGA) in the PAM sequence (the three nucleotides found straight downstream the protospacer sequence) leading to CRISPR-Cas9 mistargeting (***Supplementary figure 5A***). For pCTX15, *bla*$_{CTX\text{-}M\text{-}15}$ was amplified with Phusion High-Fidelity polymerase (Thermo Scientific) from *E. coli* ST131 strain EC958 (3), using primers introducing either a HindIII or a SalI restriction site on the amplicon edges (***Supplementary table 3***). The amplicon was purified using Monarch® DNA Gel Extraction Kit (NEB) and cloned into the digested pSEVA251 using HindIII-HF, SalI-HF and T4 DNA ligase (all from NEB). After generating pCTX15, pCTRL was obtained by introducing the PAM mutation via site-directed mutagenesis using the Phusion Site-Directed Mutagenesis Kit (Thermo Scientific). The final sequences were confirmed by Sanger sequencing and the respective plasmids were transformed into electrocompetent *E. coli S17-1λ*pir carrying pACYCduet-1.

#### *Supplementary method 2* – CRISPR-Cas9 functionality assay

For the CRISPR-Cas9 functionality assay, liquid mating was performed between *E. coli S17-1λ*pir carrying pACYCduet-1 and either pCTX15 or pCTRL (donors) and cefotaxime-resistant or cefotaxime-sensitive escapers from Ecp1-I and Ecp1-II (recipients). pACYCduet-1 was used in the donor to confer resistance to Cm, as the assay was performed under Cm selection to ensure pKJK5::csgc[*bla*$_{CTX\text{-}M\text{-}15}$] maintenance in the escapers. Donors were grown O/N in 5 mL LB + Cm + Km. Recipients were grown O/N in 600 µL LB + Cm in static conditions, using 48 well-plates. The following day, donors were washed twice with 5 mL NaCl and OD600 adjusted to 0.5 – 0.6, recipients were diluted 1000-fold in NaCl. 50 µL of donor and 50 µL of recipient were inoculated into 600 µL LB +

Cm and incubated O/N at 50 rpm. Three replicate experiments for each donor-recipient combination were performed, alongside donor-only, recipient-only and NaCl-only controls. After incubation, selective plating of serial dilutions was used to quantify the proportion of 1) donors (Cm + Km), 2) recipients (Gm + Cm) and 3) transconjugants (*E. coli* ST131 escapers carrying either pCTX15 or pCTRL, Gm + Cm + Km). For each isolate, a cefotaxime-resistant escaper with a genotypically characterized deletion in the csgc (C_D) and a cefotaxime-resistant escaper with no amplification of csgc (C_NA) were used as negative controls together with a transconjugant from the [NT] treatment (C_NT), which was used as a non-targeting control. For each isolate, two independent experiments were performed to cover all the escapers. The same controls were used for all experiments within an isolate. Conjugation efficiencies were obtained for each plasmid and the relative conjugation efficiencies (pCTRL/pCTX15) were calculated for each sample by dividing the pCTRL conjugation efficiency of each replicate by the average conjugation efficiency of pCTX15. Before performing the assay with escapers, the validity of the assay was verified by mating *E. coli S17-1λ*pir carrying either pCTX15 or pCTRL (donors) and MG1655 pKJK5::csgc[NT] or MG1655 pKJK5::csgc[*bla*$_{CTX\text{-}M\text{-}15}$] (recipients) following a similar methodology to the one described above (***Supplementary figure 5A***).

### SUPPLEMENTARY TABLES

**Supplementary table 1. Bacteria**

| Bacteria | Source | Chromosomal marker | Plasmids |
|---|---|---|---|
| Ecp1-I (ST131) | *This study* | *aacC1* between *yjjP* (*EC958_0094*) and *yjjQ* (*EC958_0095*) | |
| Ecp1-II (ST131) | *This study* | *aacC1* in *xylH* (*EC958_3974*) | |
| 26-I (ST131) | *This study* | *aacC1* in *ygjM* (*EC958_2982*) | |
| 26-II (ST131) | *This study* | None | |
| *E. coli* K-12 MG1655 | (4) | *mCherry*<br>$lacI^q$- repression of *gfp* expression | pKJK5::csgc[$bla_{CTX-M-15}$] or [NT] |
| *E. coli S17-1 λ* pir | *Common laboratory strain* | | pCTX15<br>pCTRL<br>pACYCduet-1 |
| *E. coli* DH5::SmR | (5) | *aad* | |

**Supplementary table 2. Plasmids**

| Plasmid | Payload | Source |
|---|---|---|
| pKJK5::csgc[$bla_{CTX-M-15}$] | csgc (*cas9, sgRNA, gfp, catB*)<br>Cm | This study<br>(***Supplementary method 1***) |
| pKJK5::csgc[NT] | csgc (*cas9, sgRNA, gfp, catB*)<br>Cm | This study<br>(***Supplementary method 1***) |
| pSEVA251 | Km | (2) |
| pCTX15 | Km + CTX<br>$bla_{CTX-M-15}$ inserted into pSEVA251 | This study<br>(***Supplementary method 1***) |
| pCTRL | Km + CTX<br>Mutated pCTX15; Silent point mutation in $bla_{CTX-M-15}$ | This study<br>(***Supplementary method 1***) |
| pACYCduet-1 | Cm | Novagen (Milipore) |

**Supplementary table 3. Primers.**

| ID | Oligo | DNA sequence (5' – 3') |
|---|---|---|
| | [$bla_{CTX-M-15}$] top | CTAGTCGCGTGATACCACTTCACCTGTTTTAGAGCT |
| | [$bla_{CTX-M-15}$] bottom | CTAAAACAGGTGAAGTGGTATCACGCGA |

| | | |
|---|---|---|
| | [NT] top | CTAGTGGTAAGACCATTAGAAGTAGGTTTTAGAGCT |
| | [NT] bottom | CTAAAACCTACTTCTAATGGTCTTACCA |
| 1 | CTX15_Fw | GCTGCACCGGTGGTATTGCCTT |
| 2 | CTX15_Rv | CGGCGGACGTACAGCAAAAACT |
| 3 | CTX15outside_Rv | CTGCCGATGACTATGCGCAC |
| 4 | IS*Ecp1*_Fw | CGGTGGGAAAAATGCCCTGC |
| 5 | IS*26*del_FW | CTTGCTCCTTGAGTATCGTTGCGCACCAGA |
| 6 | IS*26*del_Rv | ACCATGATGCCGACTATGGCGTGCTGAATG |
| | 16S_Fw | AGAGTTTGATCMTGGCTCAG |
| | 16S_Rv | ACCTTGTTACGACTT |
| 7 | Intl1_Fw | TAACATCAAGGCCCGATCCT |
| 8 | Cas9.1_Rv | GTCGTAGGTGTCCTTGCTCA |
| 9 | Cas9.2_Fw | TGAGCAAGGACACCTACGAC |
| 10 | Cas9.2_Rv | CGCTTCAGCTGCTTCATCAC |
| 11 | Cas9.3_Fw | GTGATGAAGCAGCTGAAGCG |
| 12 | Cas9.3_Rv | TTCACGATGTTCACCTGCGG |
| 13 | Cas9.4_Fw | CCGCAGGTGAACATCGTGAA |
| 14 | GFPbw_Rv | CTTCGCCCTTGCGCATTCTT |
| 15 | SpacersgRNA_Fw | AGCTAGCTCAGTCCTAGGTA |
| 16 | SpacersgRNA_Rv | GTTGATAACGGACTAGCCTT |
| | CmAgeI_Fw | TAGCACCGGTTTACGCACCACCCCGTCAGTAGCT |
| | CmAgeI_Rv | ATCCACCGGTAAAAATTACGCCCCGCCCTGCCA |
| | HindIIICTX15_Fw | CGTAAGCTTGCAGTACCAGCGTACGGCCC |
| | SalICTX15_Rv | GTAGTCGACGCTTATGGCCTGGTATGCGC |

### SUPPLEMENTARY FIGURES

***Supplementary figure 1:***

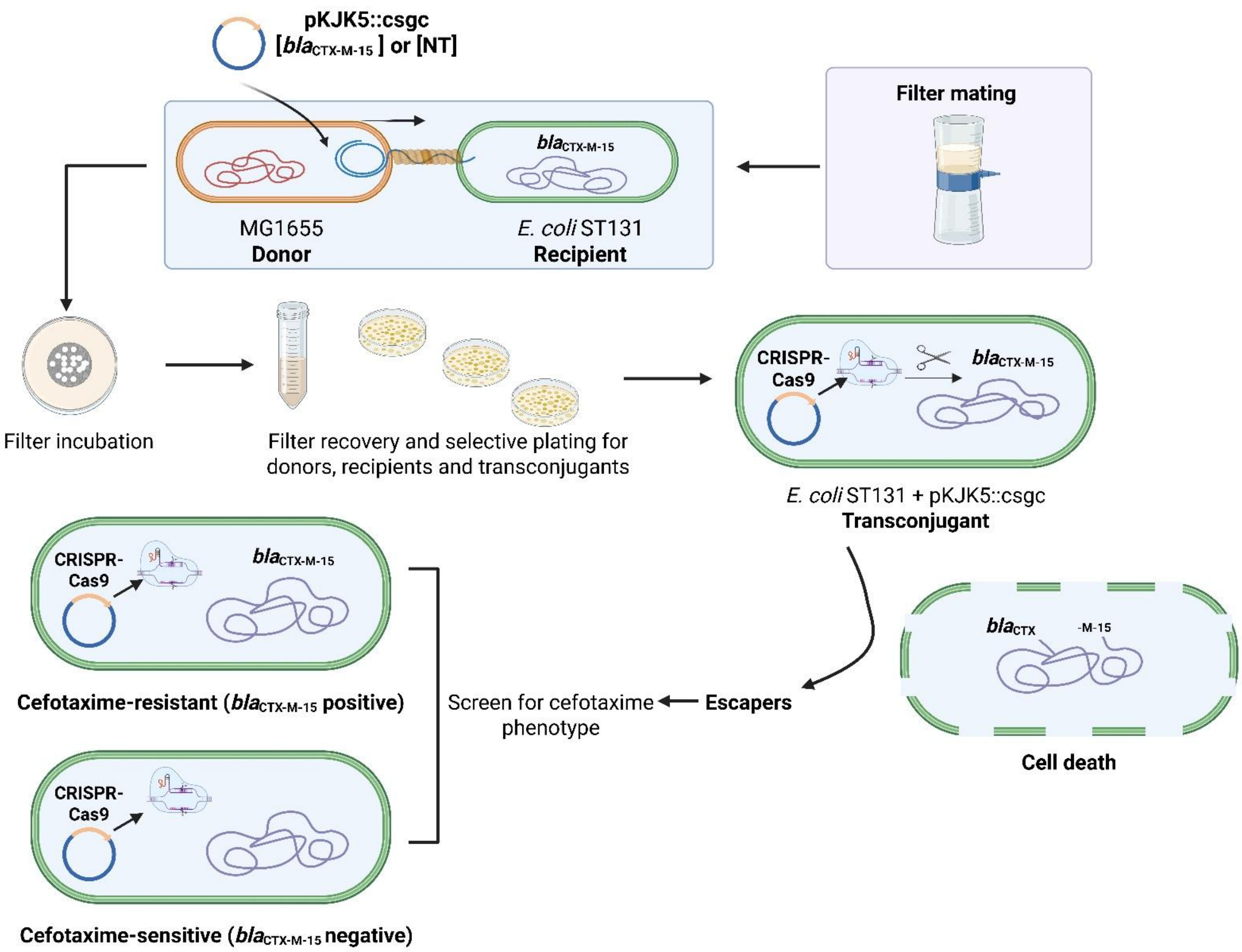


***Supplementary figure 1:*** Experimental design of the filter-mating conjugation assay to deliver pKJK5::csgc[*bla*$_{CTX-M-15}$] or [NT] from the donor *E. coli* MG1655 to the respective *E. coli* ST131 isolates. Cell death was the main CRISPR-Cas9 outcome due to *bla*$_{CTX-M-15}$ cleavage. Additionally, viable transconjugants (*E. coli* ST131 carrying pKJK5::csgc[*bla*$_{CTX-M-15}$]), also known as escapers, were found and phenotypically screened for cefotaxime, which revealed the presence of both cefotaxime-resistant (*bla*$_{CTX-M-15}$-positive) and cefotaxime-sensitive (*bla*$_{CTX-M-15}$-negative) escapers. Created in BioRender (2024).

***Supplementary figure 2*:**

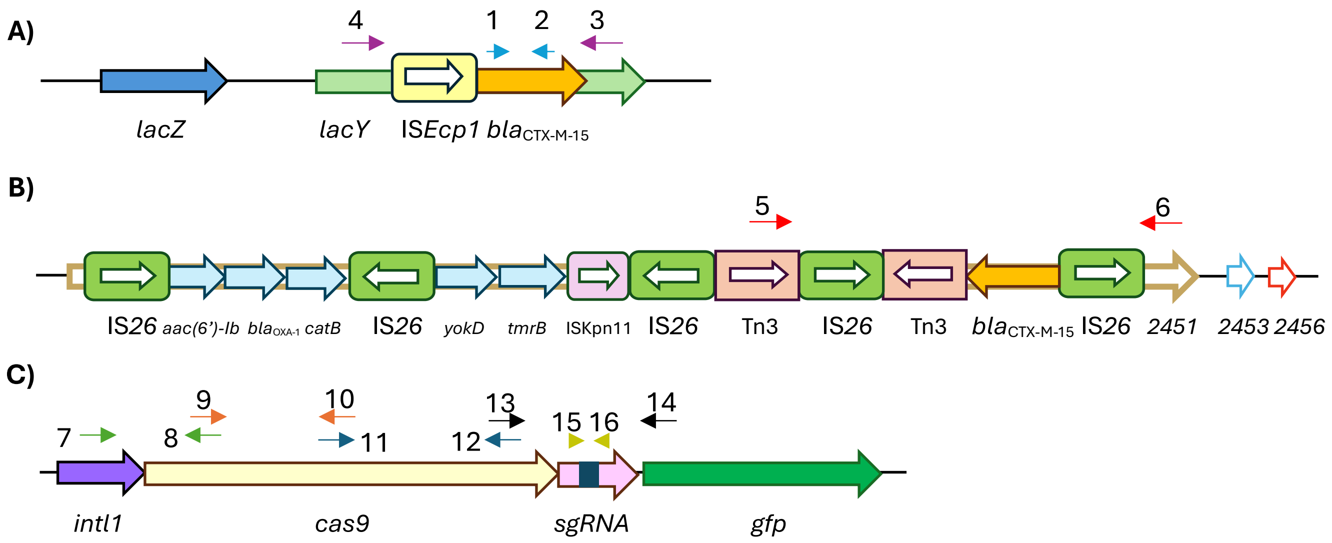


***Supplementary figure 2****:* Schematic overview of the *bla*$_{\text{CTX-M-15}}$ genetic context for isolates Ecp1-I and Ecp1-II (**A**), *bla*$_{\text{CTX-M-15}}$ genetic context for isolate 26-II (**B**) and the csgc cassette from pKJK5::csgc (**C**). ORFs and gene distances not drawn to scale. Primers used across the study can be found as sets of coloured arrows. **A**) Blue primers (1-2) show amplification of *bla*$_{\text{CTX-M-15}}$ used in cefotaxime-resistant and cefotaxime-sensitive escapers from Ecp1-I and Ecp1-II. Purple primers (3-4) show amplification of IS*Ecp1* + *bla*$_{\text{CTX-M-15}}$ used for cefotaxime-sensitive escapers from Ecp1-I and Ecp1-II. **B**) Amplification of the *bla*$_{\text{CTX-M-15}}$ PCT (5-6) **C**) Amplification of *cas9* and *sgRNA* with four sets of overlapping primers (7-8, 9-10, 11-12 and 13-14) and a fifth set covering the *sgRNA* (primers 15-16). All primers can be found in ***Supplementary table 3***. 2451, 2453 and 2456 represent the genes with unknown function *EC958_2451, EC958_2453 and EC958_2456*.

***Supplementary figure 3*:**

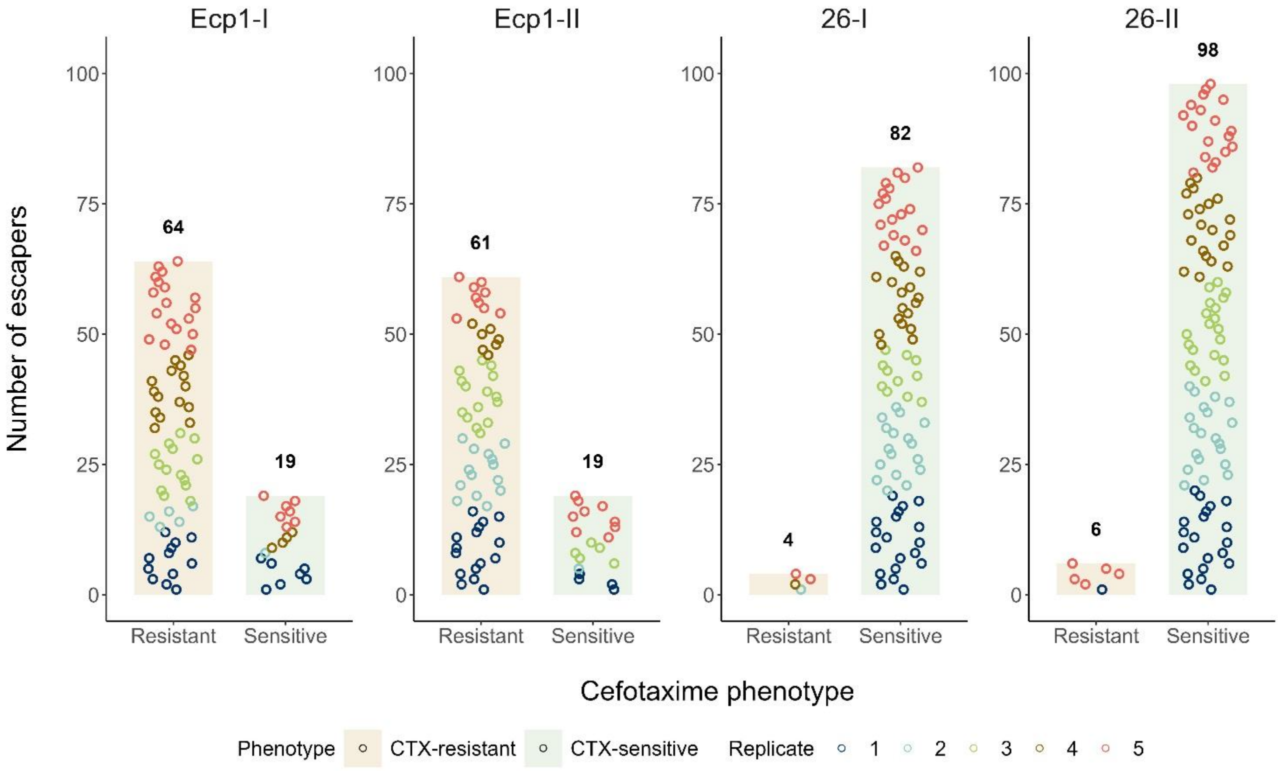


***Supplementary figure 3***: Total escapers recovered per isolate partitioned by cefotaxime phenotype and replicate. The circles represent each individual escaper, and the colour represents the filter-mating replicate it was recovered from. The total amount of escapers combines escapers directly recovered from the filter-mating assay and from the frozen stocks at a later timepoint. Only few cefotaxime-resistant escapers were recovered for isolates 26-I and 26-II due to escape frequencies being close to the limit of detection of the assay.

***Supplementary figure 4*:**

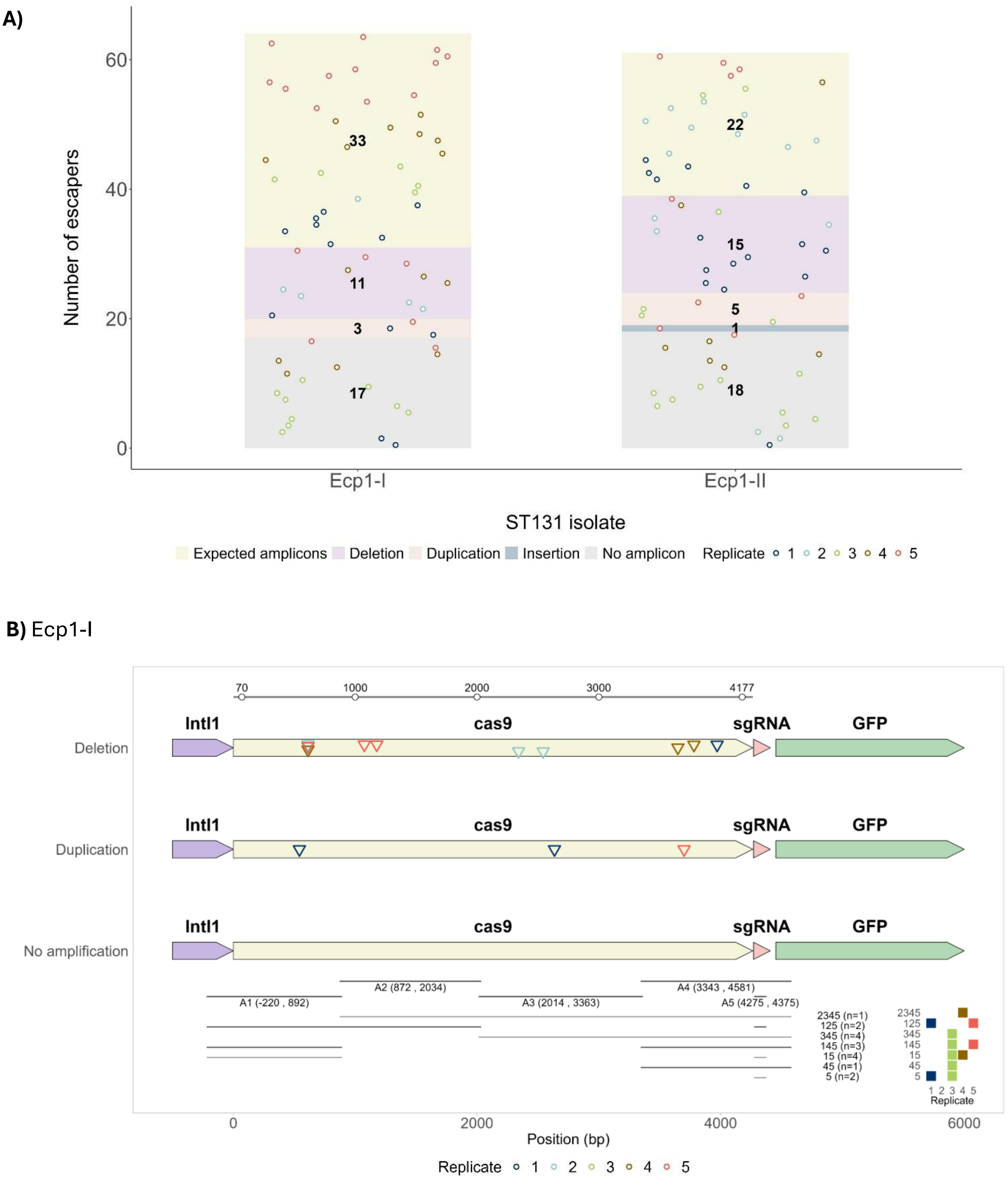

**C)** Ecp1-II

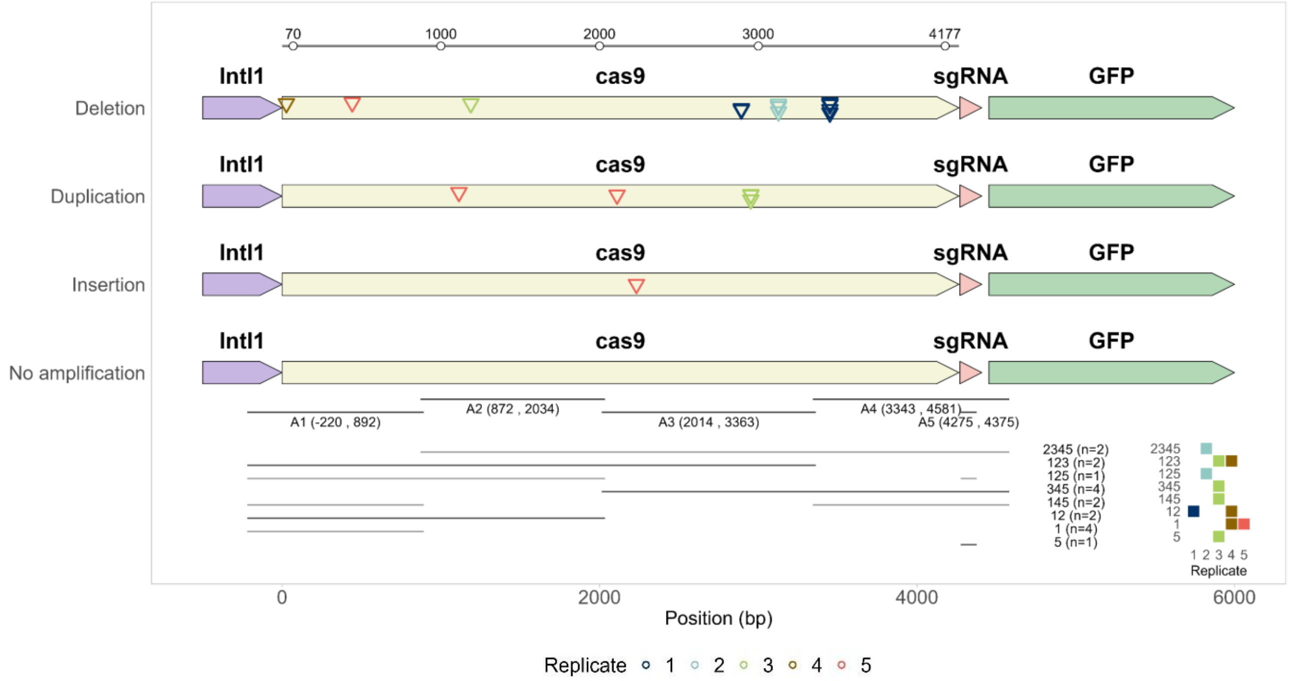


***Supplementary figure 4***: Genotypic characterization of CRISPR-Cas9 (csgc) in cefotaxime-resistant escapers from isolates Ecp1-I and Ecp1-II. **A**) Summary of results of the genotypic csgc integrity assay. The stacked bars represent the total amount of escapers distributed by expected lengths of csgc amplicons, deletions, duplications, insertions or no amplification for a single or multiple set of primers covering *cas9* and *sgRNA*. Each circle represents an escaper, and the colour represents the filter-mating replicate it was recovered from. **B-C**) Genetic map of mutations identified across the csgc in cefotaxime-resistant escapers from isolates Ecp1-I (**B**) and Ecp1-II (**C**). Triangles denote the initial position within the *cas9* gene where mutations were detected. The “No amplification” genetic map shows the five expected amplicons covering the csgc (A1 to A5), and below, the distinct csgc genotypes observed in escapers lacking one or multiple amplicons are represented along with the corresponding abundances. The table in the lower right corner indicates the original replicate where these escapers were recovered from. The ruler at the top of the figure covers the *cas9* module (0 to 4264 bp), allowing visualization of both the promoter region (7 to 69 bp) and the Cas9 coding sequence (70 to 4177 bp).

***Supplementary figure 5***:

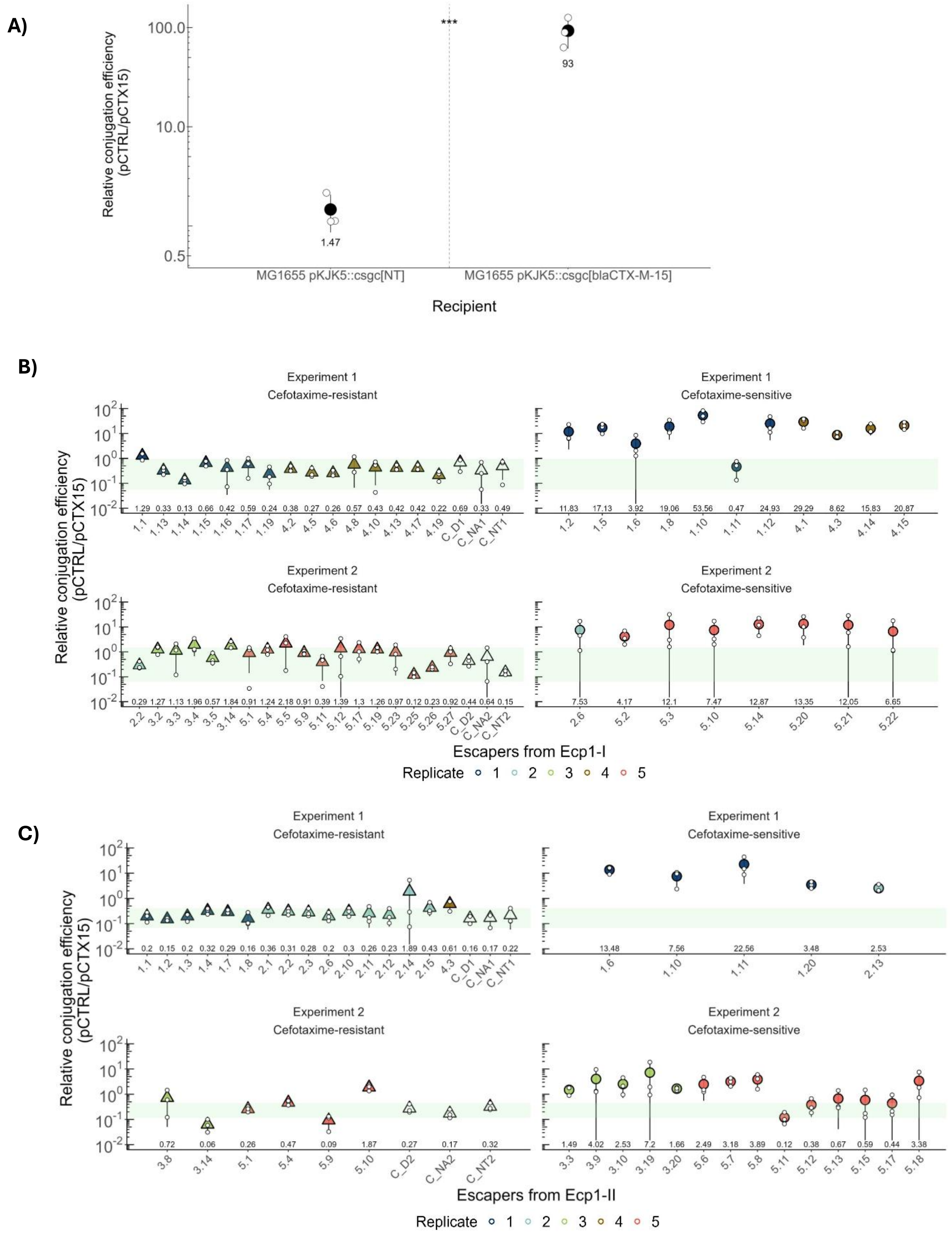

***Supplementary figure 5***: Phenotypic CRISPR-Cas9 functionality assay showing that functional pKJK5::csgc[$bla_{\text{CTX-M-15}}$] protects against acquisition of plasmid pCTX15 in comparison with pCTRL. **A**) Assay validation. The relative conjugation efficiencies (pCTRL/pCTX15) when mating either *E. coli S17-1* $\lambda$ pir pCTX15 or pCTRL (donors) with either MG1655 pKJK5::csgc[$bla_{\text{CTX-M-15}}$] or MG1655 pKJK5::csgc[NT] (recipients). Relative conjugation efficiencies were calculated by dividing the pCTRL conjugation efficiency of each replicate by the average pCTX15 conjugation efficiency. High values translate to higher conjugation efficiency for pCTRL than pCTX15 which indicates protection against pCTX15 acquisition. In contrast, low values represent similar or lower conjugation efficiency of pCTRL compared to pCTX15 which indicates lack of protection against pCTX15 acquisition. Significantly higher relative conjugation efficiency (pCTRL/pCTX15) was found for *E. coli* MG1655 pKJK5::csgc[$bla_{\text{CTX-M-15}}$] compared to the non-targeting CRISPR-Cas9 system (*E. coli* MG1655 pKJK5::csgc[NT]) ($p < 0.001$). The means and standard deviation are represented as a circle and bars. The relative conjugation efficiencies for each pCTRL replicate are represented with white circles. **B)** Escapers from isolate Ecp1-I. **C**) Escapers from isolate Ecp1-II. For both isolates, cefotaxime-resistant escapers showed significantly lower pCTR/pCTX15 ($p < 0.001$) than cefotaxime-sensitive escapers, indicating dysfunctional CRISPR-Cas9 systems. C_D, C_NA and C_NT are controls for non-functional and non-targeting CRISPR-Cas9 (see ***Supplementary method 2***). The green zone covers the range of individual replicates of negative (C_D, C_NA) and non-targeting (C_NT) controls. The mean relative conjugation efficiency (pCTR/pCTX15) for each escaper is plotted as a triangle or circle and the colour represents the filter-mating replicate it was recovered from. The value of the mean is written below. The relative conjugation efficiencies for each pCTRL replicate are represented as white circles. When no transconjugants were recovered for one replicate, the limit of detection (set to 50% of detection limit which is 1 CFU for the non-diluted sample) was used. Two sets of experiments were performed to cover all escapers from an isolate. The number after the name of the controls, is indicative of the experiment. p-values <0.001 (***).

***Supplementary figure 6:***

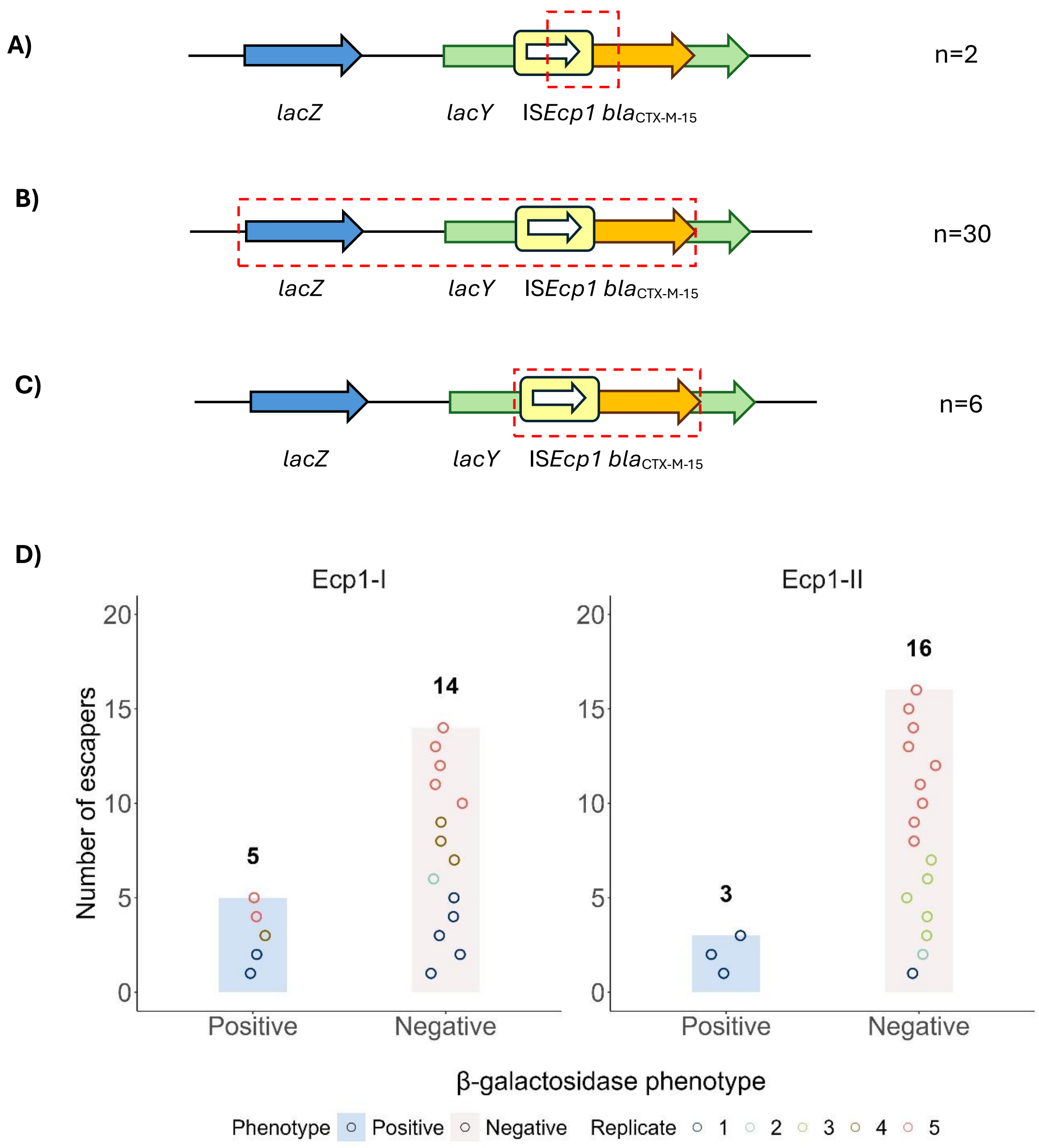


***Supplementary figure 6***: Genotypic and phenotypic profile of cefotaxime-sensitive escapers from isolates Ecp1-I and Ecp1-II. ORFs and gene distances not drawn to scale. The dashed red box represents the minimal *bla*$_{\text{CTX-M-15}}$ deletion that could be characterized combining data from both a PCR that covered IS*Ecp1* + *bla*$_{\text{CTX-M-15}}$ and the phenotypic β-galactosidase assay. **A**) β-galactosidase-positive escapers with deletions within IS*Ecp1* and *bla*$_{\text{CTX-M-15}}$, involving the loss of the protospacer (n=2). **B**) β-galactosidase-negative escapers with deletions involving at least *lacZ,* IS*Ecp1* and *bla*$_{\text{CTX-M-15}}$ (n=32). **C**) β-galactosidase-positive escapers with deletions involving at least IS*Ecp1* and *bla*$_{\text{CTX-M-15}}$ (n=6) and starting somewhere downstream *lacZ*. **D**) Results of the phenotypic β-galactosidase assay. The circles represent each escaper, and colour represents the filter-mating replicate it was recovered from.

***Supplementary figure 7:***

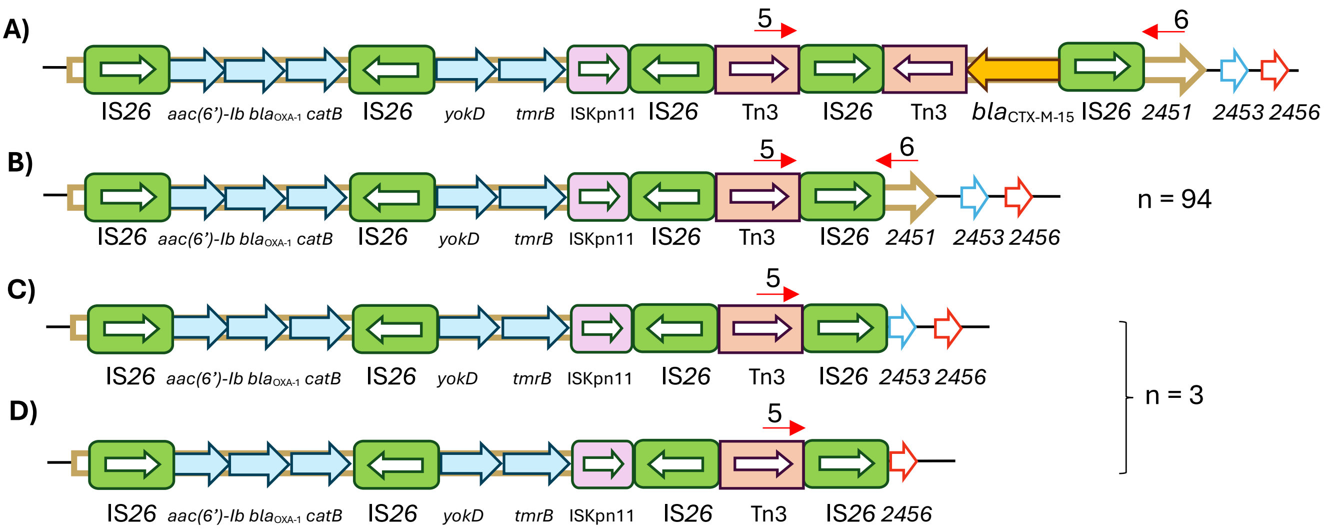


***Supplementary figure 7:*** Genotypes identified from long-read WGS data for the cefotaxime-sensitive escapers from isolate 26-II. ORFs and gene distances not drawn to scale. **A**) Genetic context of *bla*$_{CTX-M-15}$ in the ancestral isolate. The primers used to amplify the *bla*$_{CTX-M-15}$ PCT are shown as red arrows (5-6). **B**) Deletion of the *bla*$_{CTX-M-15}$ PCT, observed for 8/10 escapers in WGS and a total of 94 out of 98 with PCR. **C**) and **D**) show larger deletions including the *bla*$_{CTX-M-15}$ PCT with downstream chromosomal sequence. Each of the deletions were found for 1 escaper in the WGS and a third escaper showed a PCR result compatible with this type of deletions. 2451, 2453 and 2456 represent the genes with unknown function *EC958_2451, EC958_2453 and EC958_2456*.

***Supplementary figure 8***:

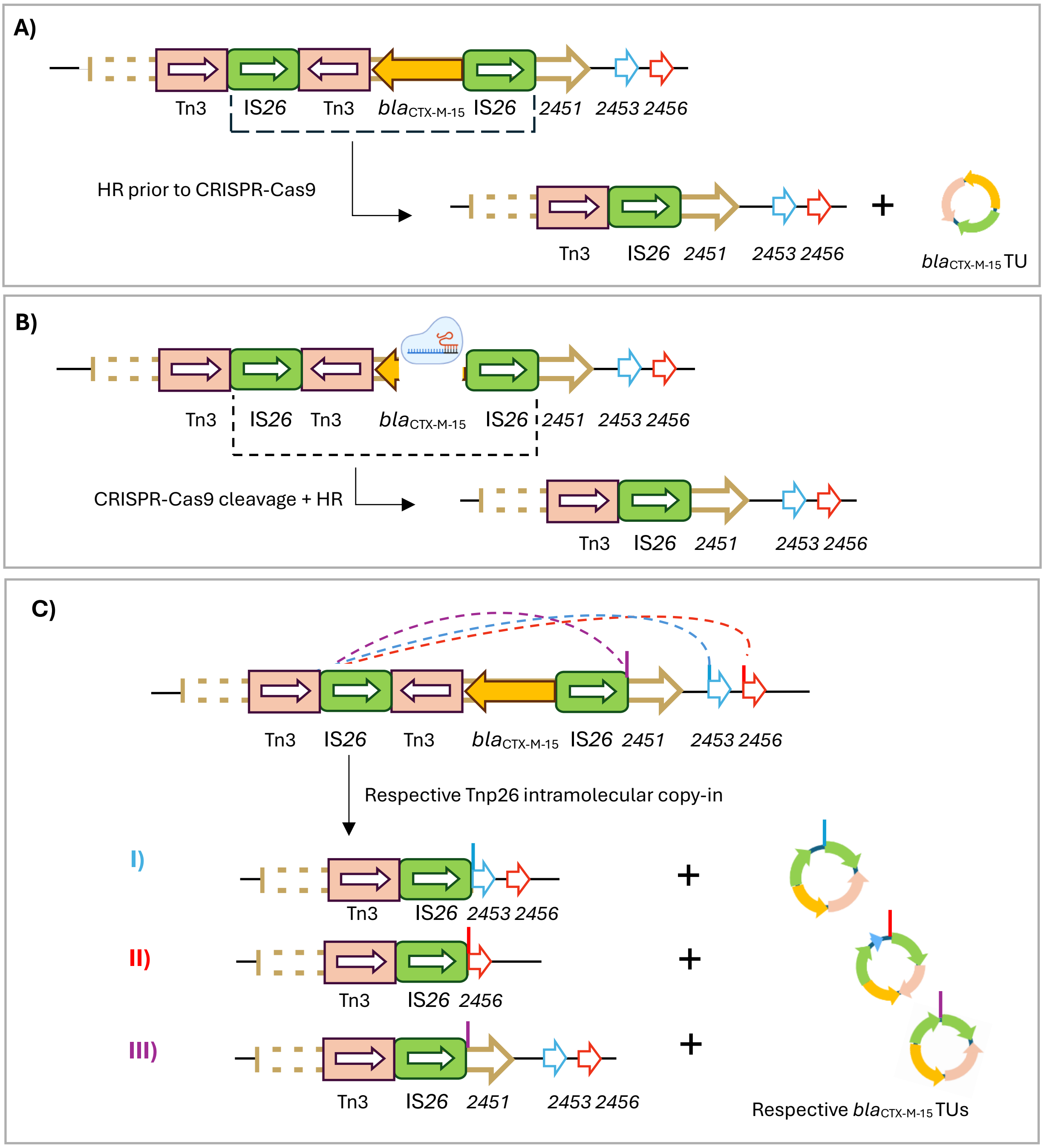


***Supplementary figure 8:*** Models explaining the potential mechanisms leading to the *bla*$_{\text{CTX-M-15}}$ deletions found in cefotaxime-sensitive escapers from isolate 26-II. ORFs and gene distances not drawn to scale. Only the relevant sequence containing the *bla*$_{\text{CTX-M-15}}$ PCT is shown from the 5-copy IS*26* sequence. **A**) Loss of *bla*$_{\text{CTX-M-15}}$ through homologous recombination (HR) prior to CRISPR-Cas9 treatment, which leads to cell survival due to absence of the target gene in the chromosome. The excised *bla*$_{\text{CTX-M-15}}$ TU could be spontaneously lost or targeted by CRISPR-Cas9. **B**) Loss of *bla*$_{\text{CTX-M-15}}$ after CRISPR-Cas9 cleavage through HR between the two surrounding IS*26*. **C**) Loss of *bla*$_{\text{CTX-M-15}}$ through a Tnp26-dependent intramolecular copy-in mechanism using the respective target sites in the chromosome (I, II and III; represented as blue, red or purple vertical lines) and generating the respective TU molecules, that could be spontaneously lost or targeted by CRISPR-Cas9. 2451, 2453 and 2456 represent the genes with unknown function *EC958_2451, EC958_2453 and EC958_2456*.

***Supplementary References***


1. Sünderhauf D, Klümper U, Pursey E, Westra ER, Gaze WH, van Houte S. Removal of AMR plasmids using a mobile, broad host-range CRISPR-Cas9 delivery tool. Microbiology (Reading). 2023 May;169(5):001334.

2. Silva-Rocha R, Martínez-García E, Calles B, Chavarría M, Arce-Rodríguez A, De Las Heras A, et al. The Standard European Vector Architecture (SEVA): a coherent platform for the analysis and deployment of complex prokaryotic phenotypes. Nucleic Acids Res. 2013 Jan 1;41(Database issue):D666-75.

3. Forde BM, Ben Zakour NL, Stanton-Cook M, Phan MD, Totsika M, Peters KM, et al. The Complete Genome Sequence of Escherichia coli EC958: A High Quality Reference Sequence for the Globally Disseminated Multidrug Resistant E. coli O25b:H4-ST131 Clone. PLoS One. 2014 Aug 15;9(8):e104400.

4. Klümper U, Riber L, Dechesne A, Sannazzarro A, Hansen LH, Sørensen SJ, et al. Broad host range plasmids can invade an unexpectedly diverse fraction of a soil bacterial community. ISME J. 2015 Mar 17;9(4):934–45.

5. Sünderhauf D, Ringger JR, Payne LJ, Pinilla-Redondo R, Gaze WH, Brown SP, et al. CRISPR-Cas is beneficial in plasmid competition, but limited by competitor toxin-antitoxin activity when horizontally transferred. bioRxiv. 2025 May 9;
